# *Mlh1* haploinsufficiency induces microsatellite instability specifically in intestine

**DOI:** 10.1101/652198

**Authors:** Kul S. Shrestha, Elli-Mari Aska, Minna M. Tuominen, Liisa Kauppi

**Affiliations:** Systems Oncology (ONCOSYS) Research Program, Research Programs Unit, Faculty of Medicine, University of Helsinki, Helsinki, Finland; Department of Biochemistry and Developmental Biology, Faculty of Medicine, University of Helsinki, Helsinki, Finland

**Keywords:** Lynch syndrome, *Mlh1* haploinsufficiency, microsatellite instability, soma-wide *Mlh1* promoter methylation

## Abstract

Tumors of Lynch syndrome (LS) patients display high levels of microsatellite instability (MSI), which results from complete loss of DNA mismatch repair (MMR), in line with Knudson’s two-hit hypothesis. Why some organs, in particular those of the gastrointestinal (GI) tract, are especially prone to tumorigenesis in LS remains unknown. We hypothesized that MMR is haploinsufficient in certain tissues, compromising microsatellite stability in a tissue-specific manner before tumorigenesis. Using mouse genetics, we tested how levels of MLH1, a central MMR protein, affect microsatellite stability *in vivo* and whether elevated MSI is detectable prior to loss of MMR function and to neoplastic growth. We assayed MSI by sensitive single-molecule PCR in normal jejunum and spleen of 4- and 12-month old *Mlh1*^*+/+*^, *Mlh1*^*+/−*^ and *Mlh1*^*−/−*^ mice, accompanied by measurements of Mlh1 mRNA and MLH1 protein expression levels.

While spleen MLH1 levels of *Mlh1*^*+/−*^ mice were, as expected, approximately 50% compared to wildtype mice, MLH1 levels in jejunum varied substantially between individual *Mlh1*^*+/−*^ mice and decreased with age. Apparently, *Mlh1*^*+/−*^ mice with soma-wide *Mlh1* promoter methylation were the most venerable to MLH1 expression level decrease in jejunum. MLH1 levels (prior to complete loss of the protein) inversely correlated with MSI severity in *Mlh1*^*+/−*^ jejunum, while in spleens of the same mice, MLH1 levels and microsatellites remained stable. Thus, *Mlh1* haploinsufficiency affects specifically the intestine where MMR levels are particularly labile, inducing MSI in normal cells long before neoplasia. A similar mechanism likely also operates in the human GI epithelium, and could explain the wide range in age of onset of LS-associated tumorigenesis.

## Introduction

Lynch syndrome (LS) is an autosomal-dominant cancer syndrome characterized by high risk of developing colorectal cancer (CRC), endometrial cancer and various other cancers. LS accounts for 1-5% of all CRC cases; individuals with LS have >80% lifetime risk of developing CRC, and are diagnosed with CRC at an average age of 44 years. LS patients carry germline heterozygous mutations in one of the DNA mismatch repair (MMR) genes, typically *MLH1* and *MSH2* [1-3]. MMR is required for the repair of single base pair mismatches and small insertion-deletion (indel) mutations arising from strand-slippage DNA replication errors [4, 5]. Microsatellites (short DNA tandem repeats), because of their repetitive nature, are highly prone to such replication errors [4, 6] leading to unrepaired indel mutations in MMR-deficient cells, a phenotype known as microsatellite instability (MSI) [4, 7]. High-level MSI is a molecular hallmark of LS patients’ tumors and reflects defective MMR [8, 9]. The conventional notion of MSI in LS-associated CRC is that it follows Knudson’s two-hit hypothesis [10]: both MMR alleles must be defective in order to trigger MSI. In this model, the first hit is inherited as a germline defect in an MMR gene, and the remaining functional MMR allele is lost (i.e. second hit) by somatic mutations, epigenetic inactivation or loss of heterozygosity (LOH), leading to complete loss of MMR function (MMR deficiency) that initiates the MSI mutator phenotype, subsequently instigating tumorigenesis [11-13].

Certain observations in LS patients, however, suggest that MMR genes may be haploinsufficient, i.e. that loss of function of just one allele may lead to loss of genome integrity. LS patients show low-level of MSI in peripheral blood leukocytes, and in normal colon mucosa [14-16]. Additionally, cell-free extracts from *Mlh1*^*+/−*^ mouse embryonic fibroblasts show decreased MMR activity [7], low MMR levels reduce MMR efficiency in non-neoplastic human cell lines [17, 18], and *Mlh1* hemizygosity increases indel mutations in cancer cell lines [19]. Given these hints of MMR haploinsufficiency, we hypothesized that reduced MMR protein levels, may provoke microsatellite stability in certain tissues, such as the highly proliferating (and tumor-prone) intestinal epithelium in mice, already prior to the “second hit”.

The MSI mutator phenotype and MMR deficiency associated tumor spectrum has been extensively characterized in constitutional *Mlh1* knock-out mice (*Mlh1*^*−/−*^ mice). These animals have a high incidence of MMR-deficient MSI-high GI tumors [7, 20-22]. Unlike in humans, GI tumorigenesis in MMR-deficient mice preferentially affects the small intestine (in particular jejunum and ileum) and not the colon [20], thus in MMR mouse models, small intestine is used to study the cellular and molecular phenotypes related to GI tumorigenesis. Here, we used *Mlh1* heterozygous mice (*Mlh1*^*+/−*^ mice) [7, 22] as model of LS to test tissue-specific MMR haploinsufficiency. As in LS, *Mlh1*^*+/−*^ mice have early-onset of GI tract tumors, and increased mortality [21]. To assess putative MMR haploinsufficiency, we determined MSI and MLH1 levels in primary cells derived from different organs, focusing on the comparison between the small intestine (highly proliferative [23] and tumor-prone) and spleen (less proliferating [24] and no MMR-dependent tumors reported). Additionally, we assayed *Mlh1* promoter methylation and tested for LOH to obtain mechanistic insights to MLH1 protein expression regulation. Further, we investigated the effect of MLH1 protein levels in tissue-specific expression of other MMR proteins.

## Methods

### Mice and genotyping

*Mlh1* mice (B6.129-*Mlh1*^tm1Rak^, strain 01XA2, National Institutes of Health, Mouse Repository, NCI-Frederick) [7] were bred and maintained according to national and institutional guidelines (Animal Experiment Board in Finland and Laboratory Animal Centre of the University of Helsinki). *Mlh1* genotyping was performed using ear pieces. DNA lysate was prepared by incubating the ear pieces overnight in 100 μl of lysis buffer supplemented with 20 μg proteinase K. The next day, lysate was boiled for 10 minutes and spun down for 1 minute at 14,000 rpm at room temperature. 0.5 μl of DNA lysate was used for genotyping. Genotyping PCR was performed using Platinum green hot start PCR master mix (Invitrogen, Carlsbad, CA). Lysis buffer components, PCR program and primers [25] used for genotyping are listed in **Supplementary table 1**. In total 58 mice (39 males and 19 females) were used in this study, the number of mice for each experiment along with the genotype and the age is indicated in the respective figures in the results section.

### Tissue collection for DNA, RNA and protein analysis

The small intestine was flushed with cold phosphate-buffered saline (PBS), cut open longitudinally and visually inspected for visible tumors. A 3 cm long piece of jejunum was cut approximately 15 cm from the pyloric sphincter. This tissue piece, henceforth referred to simply as “jejunum”, was approximately the center of the small intestine. Jejunum was inspected for any macroscopic tumor-like outgrowth under a stereoscope, and only normal-looking tissue pieces were used for the experiments. The jejunum was snap-frozen and stored at – 80°C until further use. Other tissues and *Mlh1*^*−/−*^ intestinal tumors were also collected, snap-frozen and stored at – 80°C until further use. Frozen tissues were used for subsequent DNA, RNA and protein analysis. DNA and RNA were extracted simultaneously from the same tissue piece, and protein was extracted from an adjacent tissue piece.

### DNA extraction and single-molecule MSI analysis by PCR

DNA was extracted using AllPrep DNA/RNA/Protein Mini Kit (Qiagen, Hilden, Germany) according to manufacturer’s instructions. Approximately 5 mg of tissue was used for DNA extraction. The extracted DNA was quantified using a Qubit fluorometer (Thermo Fisher Scientific, Waltham, MA) and diluted to 30 pg/μl concentration in 5 mM Tris-HCl (pH 7.5) supplemented with 5 ng/µl carrier herring sperm (Thermo Fisher Scientific). The MSI assay was performed using single-molecule PCR (SM-PCR) [26-28]. Three microsatellites, two mononucleotide repeats A27 and A33 [29], and a dinucleotide repeat D14Mit15 [7] were assayed for MSI. To ensure that individual PCRs are seeded with a single amplifiable DNA molecule, we estimated the number of amplifiable molecules by using a dilution series (30 pg, 6 pg, 4.5 pg and 0.6 pg of input DNA) for each DNA sample analyzed, similarly to previous report [30]. We determined the DNA concentration that yielded a PCR product in approximately 50% of reactions. By Poisson approximation, this PCR success rate equates to approximately one amplifiable molecule per positive reaction [27, 30, 31]. The SM-PCR programs and primers [7, 29] are listed in **Supplementary table 1**. D14Mit15 and A33 were assayed in the same PCR reaction, and a separate PCR was run for A27. Q5 High-Fidelity DNA Polymerase system (New England Biolabs, Ipswich, MA), supplemented with 1 ng/µl carrier herring sperm, was used for SM-PCR. PCR were performed in a 10 µl reaction volume. Fragment analysis was performed by combining 1μl of each PCR product, with an internal size standard (GeneScan™ 500 LIZ™ dye Size Standard, Applied Biosystems, Waltham, MA) on an ABI3730xl DNA Analyzer (Thermo Fisher Scientific). Between 116 and 205 amplifiable DNA molecules per tissue per mouse were assayed for MSI. Fragman R package was used for visualizing the electropherograms and for mutant scoring [32]. We used stringent mutant scoring criteria, adopted from [27, 33], and thus the mutation rates reported are likely a conservative estimate. Criteria for scoring microsatellite mutations were as follows:

I. A true microsatellite signal should have lower-intensity stutter peaks. Stutter peaks should display the expected size difference (that is, 1 base for mononucleotide repeats, and 2 bases for dinucleotide repeats) from the dominant peak. Reactions with peaks without stutter were considered artifacts.
II. In order for an allele to be considered as mutant, both the highest peak and the stutter peaks should shift as a single unit. Shift of the highest peak alone was not scored as a mutant.
III. If a wildtype and (apparently) mutant allele co-occurred in a single PCR, the reaction was scored as wildtype. Non-wildtype peaks were presumed to result from replication slippage during the early rounds of PCR, and thus considered artifacts.

For each microsatellite, MSI was scored separately for insertions and deletions in terms of number of single repeat-unit shifts observed.

MSI rate = total no. of single repeat-unit shifts observed / total DNA molecules analyzed

### RNA extraction and RNA expression analysis

RNA was extracted simultaneously from the same tissue pieces used for DNA extraction using AllPrep DNA/RNA/Protein Mini Kit (Qiagen) according to manufacturer’s instructions. Extracted RNA was measured using NanoDrop 1000 spectrophotometer (Thermo Fisher Scientific, Waltham, MA). 500 ng of extracted RNA was reverse-transcribed using SuperScript VILO cDNA Synthesis Kit (Invitrogen). Quantitative PCR (qPCR) was performed in CFX96 Touch Real-Time PCR Detection System (Bio-Rad) using SsoAdvance Universal SYBR Green Supermix system (Bio-Rad, Hercules, CA). Gene expression data of candidate gene was normalized to beta-actin. Data was analyzed using Bio-Rad CFX Maestro (version 1.1). mRNA expression analysis was performed for Mlh1 and Msh2. PCR programs and primers used are listed in **Supplementary table 1**.

### Western blot analysis

Frozen tissue pieces were thawed on ice, mechanically homogenized in RIPA buffer supplemented with protease inhibitor cocktail (Roche, Basel, Switzerland), incubated for 30 minutes on ice and centrifuged at 14000 rpm for 10 minutes at 4°C. The supernatant was collected and frozen until further use. Total protein concentration was measured using Pierce™ BCA™ protein assay kit (Thermo Fisher Scientific), 30 µg of total protein extract was used for western blotting. The denatured protein was run in 4–20% Mini-Protean TGX gels (Bio-Rad) and transferred to 0.2 µm nitrocellulose membrane using Trans-Blot Transfer Pack (Bio-Rad). To confirm complete protein transfer, membranes were stained with Ponceau solution for 5 minutes at room temperature. Membranes were blocked with 5% milk in PBS supplemented with 1µl/ml Tween-20 (Thermo Fisher Scientific) for 1 hour at room temperature, and incubated overnight at 4°C with primary antibodies against MLH1, and Beta-actin, followed the next day by infrared IRDye 800CW and IRDye 680RD secondary antibody incubation for 1 hour at room temperature. LI-cor Odyssey FC system (LI-COR, Nebraska, USA) was used to scan the membranes and LI-cor Image Studio lite (version 5.2) was used for image analysis. For each sample, using the same protein extract, a separate western blotting experiment was performed (as mentioned above) to study MSH2 protein expression levels using primary antibody against MSH2, with Beta-actin as loading control. Infrared IRDye 800CW and IRDye 680RD secondary antibody was used to detect MSH2 and Beta-actin, respectively. Details of antibodies and their dilutions are listed in **Supplementary table 1**. MLH1 and MSH2 protein signal intensities were normalized to Beta-actin signal intensity.

### Immunohistochemistry (IHC) and histological image analysis

The small intestine was flushed with cold 1xPBS, fixed overnight in 4% paraformaldehyde, cut open longitudinally, embedded into paraffin blocks as a “Swiss roll” [34], and sectioned at 4µm thickness. Heat induced antigen retrieval was performed for 20 minutes using 10mM citrate buffer (pH 6). MLH1 was detected using anti-MLH1 antibody (cat. no. ab92312, clone EPR3894, Abcam, Cambridge, UK, at 1:1500 dilution), and visualized using BrightVision Poly HRP goat anti-rabbit IgG (cat. no. DPVR55HRP, ImmunoLogic, Duiven, The Netherlands). Sections were counterstained with hematoxylin-eosin. Slides were scanned using 3DHistech Panoramic 250 FLASH II digital slide scanner (3DHistech, Budapest, Hungary) at 20X magnification, and images were visualized using CaseViewer (version 2.2). For each mouse, MLH1 protein levels were quantified for five randomly selected sites in the jejunum, each site consisting of approximately ten villus-crypt units, altogether approximately 50 villus-crypt units per jejunum. Quantification was performed using IHC Profiler plugin in ImageJ [35]. IHC was also performed to study MSH2 protein levels, as described above, using anti-MSH2 antibody (cat. no. D24B5, Cell Signaling Technology, Danvers, MA, at 1:1000 dilution).

### Mlh1 LOH analysis by PCR

LOH was performed in the same extracted genomic DNA that was used for MSI analysis. LOH was tested across the *Mlh1* gene by using LOH-PCR, as well as restriction fragment length polymorphism (RFLP) assay on large PCR products spanning the beginning and end of the *Mlh1* gene (**Fig. 4A**). For LOH-PCR, 50 ng of DNA was used for PCR. PCR was performed with Platinum green hot start PCR master mix (Invitrogen) using previously published primers to assess LOH at *Mlh1* [36] (for PCR conditions and primers see **Supplementary table 1)**. The RFLP assay was performed on PCR products covering 5’ and 3’ ends of *Mlh1* gene, henceforth referred to as RFLP_region 1 and RFLP_region 2, respectively. The Phusion High-Fidelity DNA Polymerase system (Thermo Fisher Scientific) was used for the PCR step; details of the PCR program and primers used are in **Supplementary table 1**. 200 ng of PCR product of RFLP_region 1 and RFLP_region 2 were digested with *Pst*I (New England Biolabs) and *Vsp*I (Thermo Fisher Scientific) restriction enzymes, respectively, according to manufacturer’s instruction. For both LOH-PCR and RFLP analysis, *Mlh1*^*+/+*^ and *Mlh1*^*−/−*^ jejunum DNA was used as normal and knock-out allele controls, respectively. LOH-PCR and digested PCR products (in RFLP assay) were analyzed by gel electrophoresis on an ethidium bromide stained 1.5% agarose gel. DNA samples were considered to have undergone LOH if the wildtype PCR product was absent in LOH-PCR, and if they displayed the same banding pattern as *Mlh1*^*−/−*^ DNA in the RFLP assay.

**Fig. 1.**
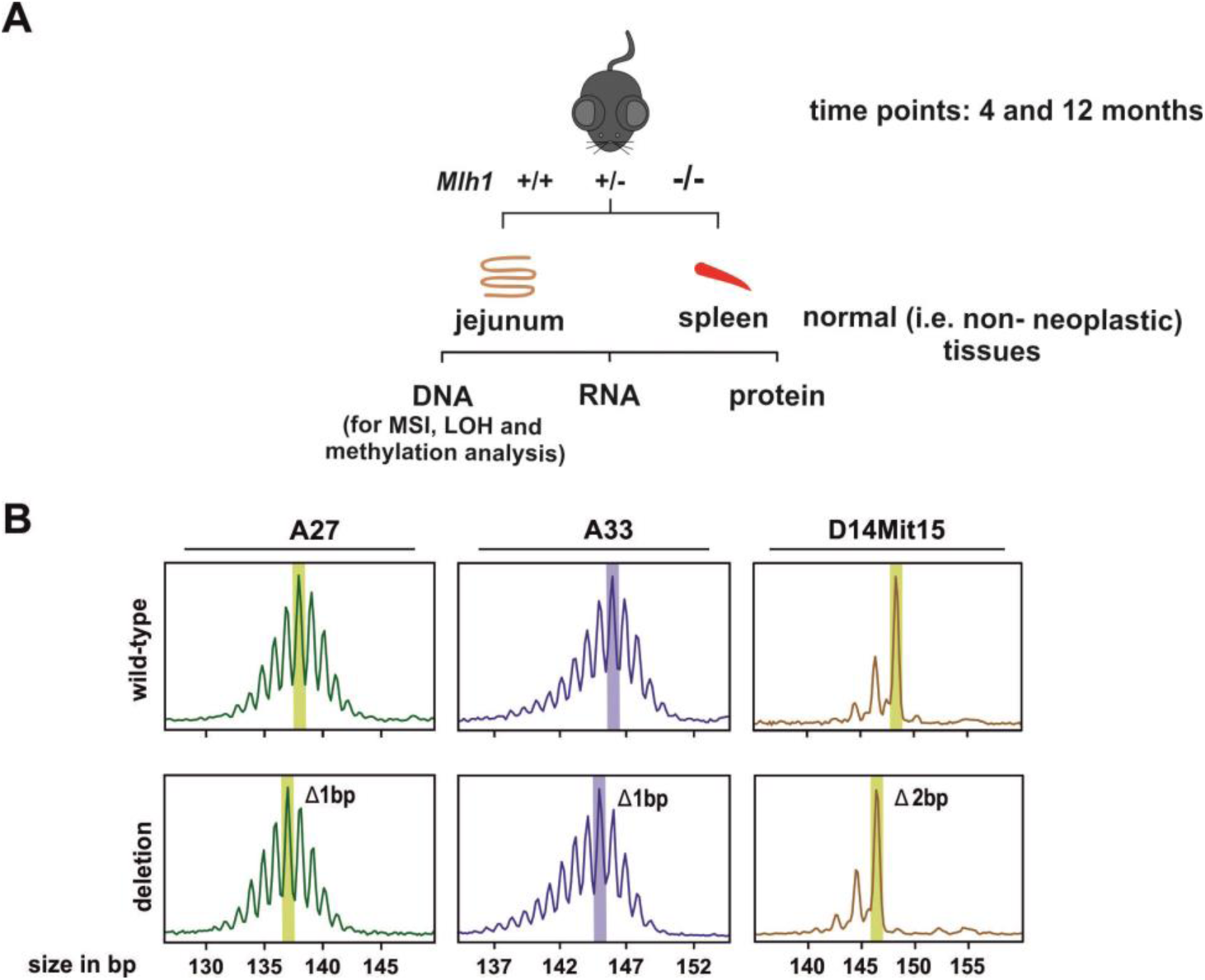
Study approach. (**A**) Workflow of the study. We tested microsatellite instability (MSI) at single-DNA molecule resolution along with analyzing *Mlh1* expression both at mRNA and protein level. In addition, we tested LOH (loss of heterozygosity) across *Mlh1* gene and assayed *Mlh1* promoter methylation status. (**B**) Representative capillary electropherograms of single-molecule PCR (SM-PCR) based MSI assay. MSI was tested at three microsatellites: two mononucleotide repeats A27 and A33, and a dinucleotide repeat D14Mit15. Top and bottom panels show electropherograms scored as wildtype and single repeat unit deletion mutant alleles, respectively. The highest peaks (indicated by shading in each electropherogram) were scored; smaller peaks flanking the highest peak are stutter peaks, a typical feature of microsatellite markers.

**Fig. 2.**
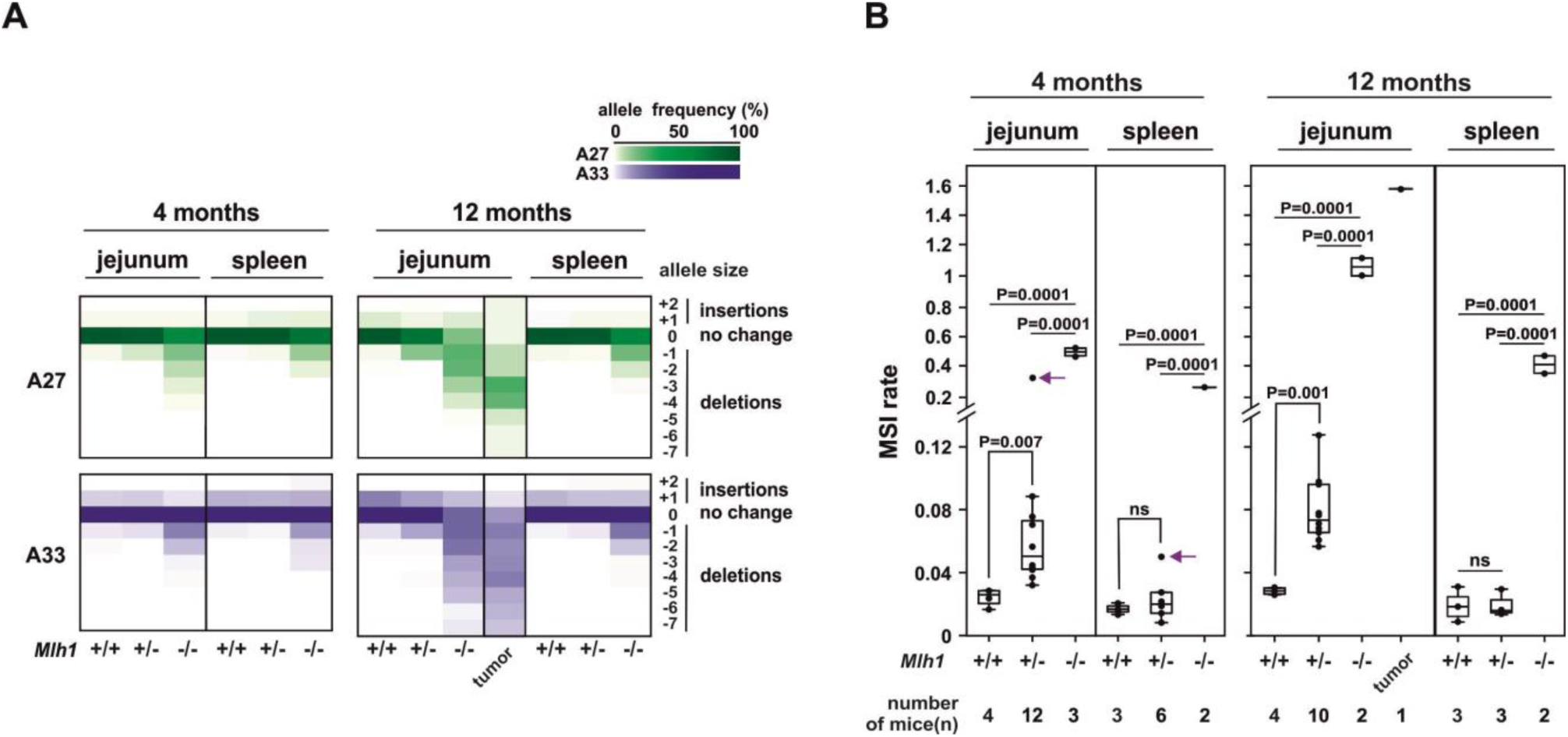
Single-molecule MSI analysis at mononucleotide repeats in jejunum and spleen at 4- and 12-month time points. (**A**) Heat map of allele frequency (%) detected for microsatellites A27 and A33. The 4-month old *Mlh1* ^*+/−*^ outlier mouse (indicated by an arrow in **Fig. 2B**) was excluded from the heat map, and is shown separately in **Supplementary fig. 3**. (**B**) MSI rate for deletions in jejunum and spleen of *Mlh1*^*+/−*^ mice. Arrow indicates the outlier *Mlh1*^*+/−*^ mouse.

**Fig. 3.**
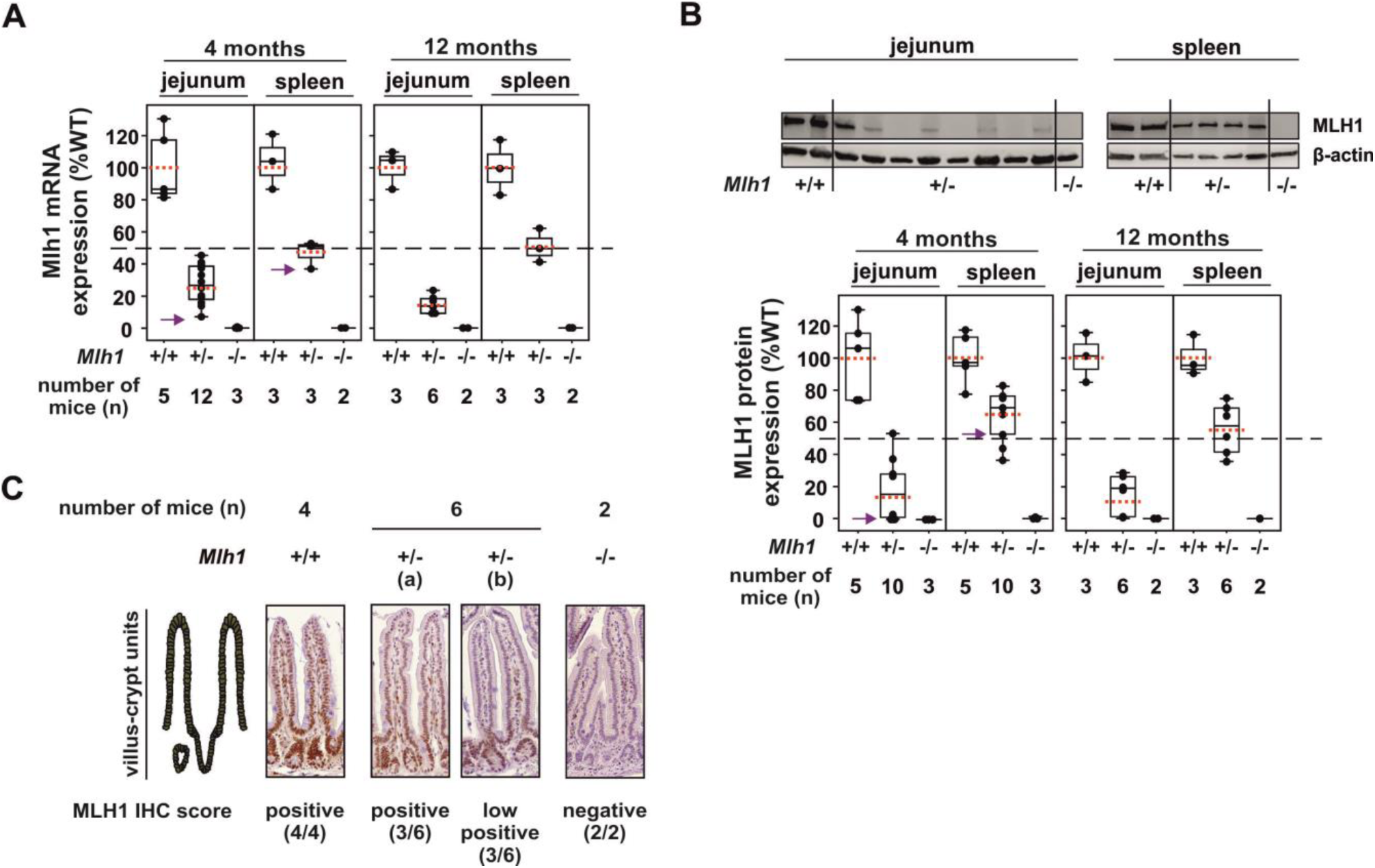
*Mlh1* gene expression analysis in jejunum and spleen of *Mlh1+/−* mice at the 4- and 12-month time points. (**A**) Mlh1 mRNA expression analysis. Arrow indicates data from the outlier *Mlh1*^*+/−*^ mouse with high deletions at mononucleotide repeats. Red dotted line across each box-plot indicates the average value. Dashed black line across the chart marks 50% expression level. (**B**) and (**C**) MLH1 protein levels analysis. (**B**) Representative image of western blot. Empty lanes in western blot do not necessarily mean absence of MLH1 but possibly MLH1 protein levels below detectable range of our western blot assay (**Supplementary fig. 7**). Boxplot shows MLH1 protein expression analysis for the western blots. The red-dotted line across each box-plot represents the average value. The dashed-horizontal line across the chart represents the 50% MLH1 protein expression. (**C**) Representative IHC images of jejunum. Middle two IHC images shows a side-by-side comparison of villus-crypt units scored as (a) positive and (b) low positive by IHC profiler.

**Fig. 4.**
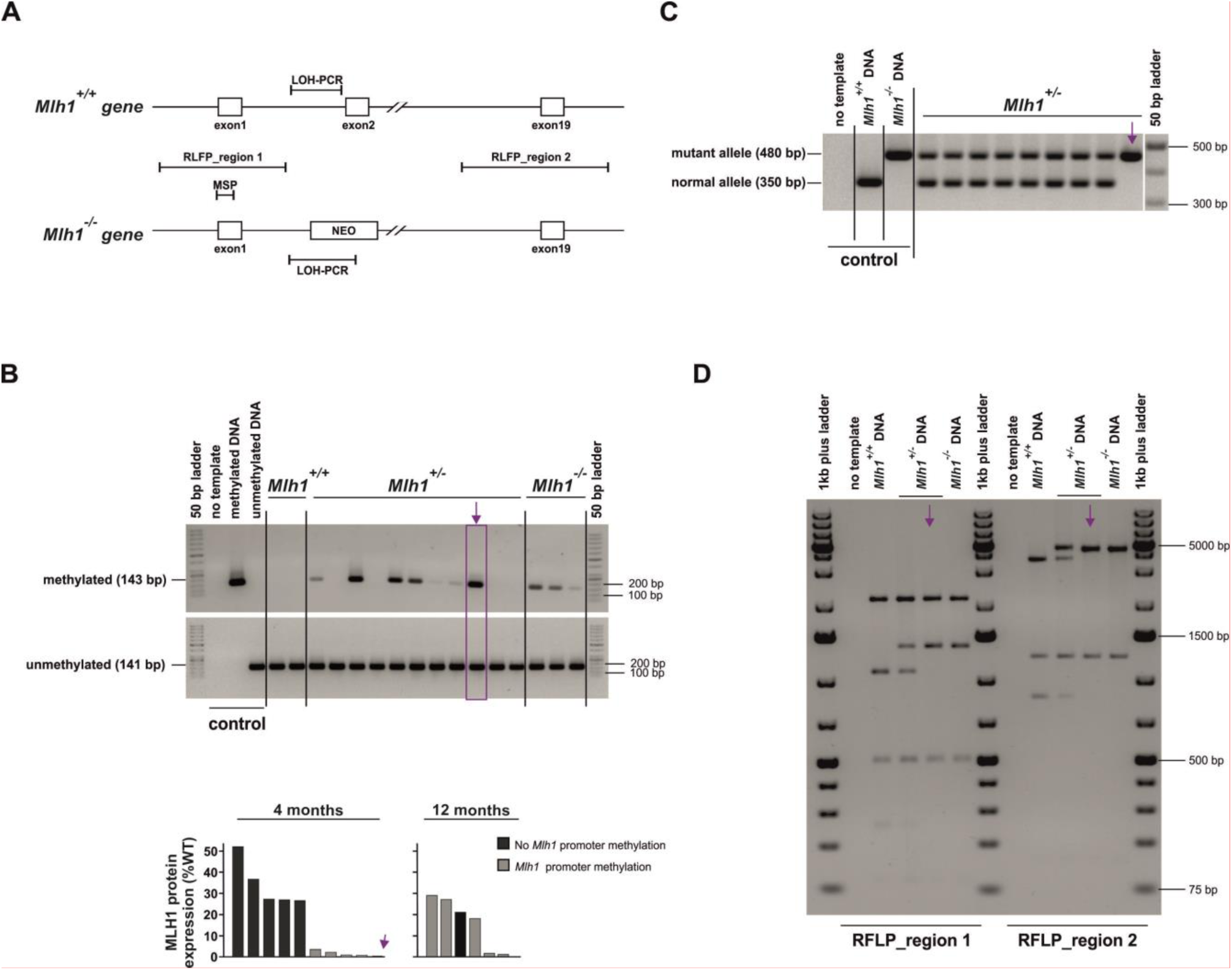
Promoter methylation and LOH assay in *Mlh1^+/−^* jejuna. (**A**) Schematic of MSP and LOH assay design. (**B)** Representative image of MSP assay (top). The bottom histogram shows co-relation between *Mlh1* promoter methylation and MLH1 protein expression, each bar represents an individual *Mlh1*^*+/−*^ mouse. MSP assay for other tissues is shown in **Supplementary fig. 5A and 5C**. Arrow indicates the outlier *Mlh1*^*+/−*^ mouse. (**C**) and (**D**) Representative images of LOH-PCR assay, and PCR-RFLP assay, respectively. Arrow indicates the outlier *Mlh1*^*+/−*^ mouse.

### Mlh1 promoter methylation analysis by methylation-specific PCR (MSP)

As for the LOH assay, we used the same extracted genomic DNA used for MSI analysis to test the *Mlh1* promoter methylation status. Methylation analysis for *Mlh1* was performed using MSP [37]. Universally methylated mouse DNA standard (cat. no. D5012, Zymo Research, Irvine, CA) and genomic DNA extracted from normal spleen was used as positive and negative controls, respectively. For MSP, 500 ng of DNA was bisulfite-converted using EZ DNA Methylation-Direct Kit (Zymo Research) according to manufacturer’s instruction. 1μl of bisulfite-converted DNA was used for PCR, which was performed using 2xZymo Taq premix. Details of the PCR program and primers [38] are in **Supplementary table 1**. PCR products were analyzed by gel electrophoresis on an ethidium bromide stained 1.5% agarose gel. Methylation status of the *Mlh1* promoter was scored qualitatively, based on presence or absence of the methylated PCR product. Further, to identity the allele (*Mlh1*^*+/+*^ or/and *Mlh1*^*−/−*^) with *Mlh1* promoter methylation in *Mlh1*^*+/−*^ tissues, we performed RFLP on 200 ng of MSP PCR-product using Hph1 (New England Biolabs) restriction enzyme accordingly to manufacture’s instruction. Universally methylated mouse DNA standard (cat. no. D5012, Zymo Research) and *Mlh1*^*−/−*^ tissues were used as *Mlh1*^*+/+*^ and *Mlh1*^*−/−*^ allele controls, respectively. Digested MSP PCR-products were run in 5% Mini-Protean precast TBE gels (Bio-Rad), post-gel run, gels were submerged into 0.5 µg/ml ethidium bromide in water for 15 minutes and imaged with Gel Doc XR+ system (Bio-Rad). Banding pattern of *Mlh1*^*+/−*^ samples were compared to the controls to identify the allele with and without *Mlh1* promoter methylation.

### Statistical analysis

Statistical testing was done using an unpaired t-test, and linear regression fitting was tested using F-test. Two-tailed P-values < 0.05 were considered statistically significant. P-values are either stated as numerical values, or indicated as follows: p-value ≥ 0.05 = not significant (ns), p-value < 0.05 = *, p-value <0.01 = **, and p-value < 0.005 = ***.

## Results

### Normal jejunum of Mlh1^*+/−*^ *mice displays MSI*

To investigate tissue- and age-specific MSI in *Mlh1*^*+/−*^ mice, we performed highly sensitive SM-PCR in jejunum and spleen of 4- and 12-month-old *Mlh1*^*+/−*^ mice (**Fig. 1A**) on three microsatellite loci (**Fig. 1B**). Mononucleotide tract A27 is an intergenic microsatellite approximately 2 kb downstream to *Epas1* gene, and A33 is an intronic microsatellite within *Epas1* gene. D14Mit15 is an intergenic microsatellite at 40 kb distance from *Ptpn20* gene.

Compared to age-matched *Mlh1*^*+/+*^ jejuna, in *Mlh1*^*+/−*^ jejuna deletions at mononucleotide repeats were elevated (observed deletions were predominantly single repeat unit, i.e. 1 bp, shifts), while the dinucleotide repeat D14Mt15 was stable (**Fig. 2A, Fig. 2B, Supplementary fig. 1A and Supplementary fig. 1B**). Compared to *Mlh1*^*+/+*^ jejuna, deletions at mononucleotide repeats in *Mlh1*^*+/−*^ jejuna were 2- and 5-fold higher, and compared to *Mlh1*^*−/−*^ jejuna 9- and 13-fold lower at 4 and 12 months, respectively (**Fig. 2B**). 12-month *Mlh1*^*+/−*^ jejuna showed 2-fold increase in deletions compared to 4-months *Mlh1*^*+/−*^ jejuna (p=0.006). Unlike *Mlh1*^*+/−*^ jejuna, *Mlh1*^*+/−*^ spleens were microsatellite stable at both time points (**Fig. 2A and Fig. 2B**).

In contrast to *Mlh1*^*+/−*^ tissues, in *Mlh1*^*−/−*^ tissues deletion mutant alleles had bigger size shifts and were more frequent at all three microsatellites at both the time points (**Fig. 2A and Supplementary fig. 1A)**. *Mlh1*^*−/−*^ jejuna showed bigger size shifts than *Mlh1*^*−/−*^ spleens; deletion size shifts further increased in *Mlh1*^*−/−*^ intestinal tumor (**Fig. 2A and Supplementary fig. 1A**). Compared to age-matched *Mlh1*^*+/+*^ tissues, deletions at mononucleotide repeats in *Mlh1*^*−/−*^ jejuna were 21- and 60-fold higher, and in *Mlh1*^*−/−*^ spleen 15- and 24-fold higher at 4- and 12-month time point, respectively (**Fig. 2B**). For comparison, the *Mlh1*^*−/−*^ intestinal tumor showed 90- and 20-fold increase in deletions compared to 12-month *Mlh1*^*+/+*^ and *Mlh1*^*+/−*^ jejuna, respectively (**Fig. 2B**). Dinucleotide repeat D14Mit15 was stable in *Mlh1*^*+/+*^ and *Mlh1*^*+/−*^ tissues at both the time-points, *Mlh1*^*−/−*^ tissues displayed MSI at D14Mit15. Compared to mononucleotide repeats, deletions at D14Mit15 were 4- and 6-fold lower in *Mlh1*^*−/−*^ jejuna and *Mlh1*^*−/−*^ spleen, respectively.

Unlike deletions, insertion frequencies at all three microsatellites were not influenced by *Mlh1* gene dosage: *Mlh1*^*+/+*^, *Mlh1*^*+/−*^, *Mlh1*^*−/−*^ mice showed minimal or no difference in insertions in jejunum and in spleen, with the exception of A33 at the 12-month time point (**Fig. 2A, Supplementary fig. 1A, Supplementary fig. 1B and Supplementary fig. 2**). At 12 months, A33 showed a decreasing trend in insertions with decreased *Mlh1* gene dosage in jejunum (**Supplementary fig. 2**). Irrespective of genotype, tissue-of-origin or age, insertions were almost exclusively single repeat unit in size (1 bp and 2 bp for mono- and dinucleotide repeats, respectively) (**Fig. 2A and Supplementary fig. 1A**).

One 4-month old *Mlh1*^*+/−*^ mouse showed substantially elevated deletions in jejunum (6- and 4-fold higher compared to other age-matched *Mlh1*^*+/−*^ mice at mononucleotide markers and at D14Mit15, respectively; indicated by an arrow in **Fig. 2B and Supplementary fig. 1B**. With Grubbs’ test, this *Mlh1*^*+/−*^ mouse was classified as an outlier (p < 0.05). A slight increase in deletions at mononucleotide repeats was also detected in the spleen of this mouse (2-fold higher than other *Mlh1*^*+/−*^ spleen), while D14Mit15 was stable (**Fig. 2B and Supplementary fig. 1B)**. In this mouse deletions consisted of predominantly one and two repeat shifts (**Supplementary fig. 3)**. Moreover, inter-individual variation in deletions at mononucleotide repeats was evident in *Mlh1*^*+/−*^ jejuna in both age groups (**Fig. 2B**).

### Mlh1^+/−^ mice show sporadic decrease in MLH1 expression levels in jejunum

To assess whether the inter-individual variation in deletion frequencies in *Mlh1*^*+/−*^ jejuna can be explained by *Mlh1* expression levels, we quantified Mlh1 mRNA and MLH1 protein levels using qPCR and western blot, respectively. *Mlh1*^*+/−*^ spleens, which were microsatellite-stable (**Fig. 2B**), were used as control tissue. To further examine tissue-specific differences in *Mlh1* expression, we analyzed Mlh1 mRNA expression in brain, kidney and liver using qPCR. We also assessed variation in MLH1 expression by immunohistochemistry (IHC) in jejunum of 4-month old mice. To study whether levels of key MMR proteins beyond MLH1 may also be altered in *Mlh1*^*+/−*^ jejuna, we quantified Msh2 mRNA and MSH2 protein levels in jejunum of 4-month old mice, with *Mlh1*^*+/−*^ spleens as tissue control. Further, we also performed IHC to assess variation in MSH2 expression in jejunum of 4-month old mice.

In *Mlh1*^*+/−*^ jejuna, average *Mlh1* expression (both at mRNA and protein levels) was less than the expected approximately 50% of *Mlh1*^*+/+*^ jejuna (**Fig. 3A and Fig. 3B**). Further, average expression was lower in the older age group, implying age-dependent MLH1 depletion. In addition, expression levels varied between individual *Mlh1*^*+/−*^ mice; mice in the 4-month old group showed more inter-individual variation than those in the 12-month old group. At 4 months, *Mlh1*^*+/−*^ jejuna showed on average 26% Mlh1 mRNA (range: 8%-46%) and 24% (range: 0%-54%) MLH1 protein expression compared to *Mlh1*^*+/+*^ levels, and at 12 months the average Mlh1 mRNA and MLH1 protein expression decreased to 15% (range: 9%-23%) and 17% (range: 1%-29%), respectively **(Fig. 3A and Fig. 3B**). These age-specific decreases in Mlh1 mRNA and MLH1 protein were not statistically significant, however.

To investigate whether younger mice show similar aberration in *Mlh1* expression levels, we analyzed Mlh1 mRNA expression levels also at 1-month time point. As the older cohorts, 1-month *Mlh1*^*+/−*^ jejuna also showed less-than-expected- and variable Mlh1 mRNA expression levels (average: 30%, range: 21%-37%) (**Supplementary fig. 4**). 12-month old *Mlh1*^*+/−*^ mice showed a significant decrease in Mlh1 mRNA levels in jejunum (p=0.0283) when compared to 1-month old *Mlh1*^*+/−*^ mice (**Supplementary fig. 4**).

Irrespective of age, we observed approximately expected 50% Mlh1 mRNA and MLH1 protein levels in all the *Mlh1*^*+/−*^ spleens (**Fig. 3A, Fig. 3B and Supplementary fig. 4)**. Jejunum of the outlier *Mlh1*^*+/−*^ mouse showed only 8% Mlh1 mRNA expression (**Fig. 3A**) and no detectable MLH1 protein (**Fig. 3B**), while its spleen had the expected levels of Mlh1 mRNA and MLH1 protein (**Fig. 3A and Fig. 3B**).

We also assayed Mlh1 mRNA expression in brain, kidney and liver of two 4-month old *Mlh1*^*+/−*^ mice that expressed below age-average Mlh1 mRNA levels in their jejunum; Mlh1 mRNA expression in these tissues were as expected (60%, 56% and 56%, respectively) (**Supplementary fig. 5D**).

Substantial inter-individual variation in MLH1 protein expression in *Mlh1*^*+/−*^ jejuna was also detectable by IHC. Based on staining intensity, jejuna of half the *Mlh1*^*+/−*^ mice (n=6) were scored as MLH1-positive and the other half was scored as low MLH1-positive by IHC profiler (**Fig. 3C**). Upon further analyzing intestinal crypts and villi separately, in MLH1 low-positive jejuna we observed decreased MLH1 staining intensity in both; moreover, all MLH1-negative cells were located in the villi **(Supplementary fig. 6)**.

We also investigated whether expression levels of MMR components beyond MLH1, namely MSH2, were affected in *Mlh1* heterozygotes. At 4 months, *Mlh1*^*+/−*^ mice showed uniform Msh2 mRNA expression levels both in jejunum and in spleen, except for one sample: jejunum of the 4-month old outlier *Mlh1*^*+/−*^ mouse, which showed low Msh2 expression (**Supplementary fig. 8A**). At the protein level, however, MSH2 showed inter-individual variation in *Mlh1*^*+/−*^ jejuna similar to MLH1. That is, *Mlh1*^*+/−*^ jejunum samples with low MLH1 levels also had low MSH2 levels (**Supplementary fig. 8B**). Additionally, MSH2 protein expression was affected by *Mlh1* dosage in both jejunum and spleen. Compared to *Mlh1*^*+/+*^ tissue, MSH2 expression was lower in *Mlh1*^*+/−*^ tissue and in *Mlh1*^*−/−*^ tissue (**Supplementary fig. 8B**). Interestingly, 6 out of 10 *Mlh1*^*+/−*^ jejuna expressed lower MSH2 than *Mlh1*^*−/−*^ jejuna. IHC was less sensitive (compared to western blot) to detect the inter-individual variation in MSH2 expression levels. In IHC analysis, based on the staining intensity, all samples irrespective of the genotype (n=4, 6 and 2 for *Mlh1*^*+/+*^, *Mlh1*^*+/−*^ and *Mlh1*^*+/−*^ jejuna, respectively) were scored as MSH2-positive expect one *Mlh1*^*+/−*^ jejuna which was scored as low MSH2-positive (**Supplementary fig. 8C**).

### Tissue-specific depletion of MLH1 in jejuna of Mlh1^+/−^ mice with Mlh1 promoter methylation

The variable MLH1 protein levels in jejunum of *Mlh1*^*+/−*^ mice ranging from approximately expected 50% of *Mlh1*^*+/+*^ to no expression (**Fig. 3B**) prompted us to investigate the cause of this sporadic MLH1 depletion. To distinguish between a genetic (loss of heterozygosity) and epigenetic mechanism (methylation of the *Mlh1* promoter), we performed LOH and MSP assays, respectively (**Fig. 4A**). In addition, we identified the *Mlh1* allele harboring *Mlh1* promoter methylation by performing RFLP in MSP PCR-products.

*Mlh1* promoter methylation in jejunum was common (detected in 7/12 and 9/10 *Mlh1*^*+/−*^ mice at 4 and 12 months, respectively; see **Fig. 4B)**. Irrespective of age, all *Mlh1*^*+/−*^ mice with extremely low (less than 3%) MLH1 protein expression in jejunum (5/10 and 2/6 *Mlh1*^*+/−*^ mice at 4 and 12 months, respectively) displayed *Mlh1* promoter methylation (**Fig. 4B**). Additionally, three of the four 12-month old *Mlh1*^*+/−*^ mice with higher level of MLH1 expression (range: 18%-29%) showed *Mlh1* promoter methylation, while none (out of six) of the 4-month old *Mlh1*^*+/−*^ mice with higher MLH1 levels (range: 18%-52%) had *Mlh1* promoter methylation (**Fig. 4B**). In addition, we also tested *Mlh1* promoter methylation in 1-month old *Mlh1*^*+/−*^ jejuna, 3 out of 5 *Mlh1*^*+/−*^ mice showed *Mlh1* promoter methylation in this cohort.

We also tested *Mlh1* promoter methylation in spleen, all the *Mlh1*^*+/−*^ mice (irrespective of age) with *Mlh1* promoter methylation in jejuna showed *Mlh1* promoter methylation in their spleen, others (i.e. *Mlh1*^*+/−*^ mice without *Mlh1* promoter methylation in jejuna) did not show *Mlh1* promoter methylation in their spleen (**Supplementary fig. 5A**). Further, performing RFLP in the MSP PCR-products, we identified the *Mlh1*^*−/−*^ allele as the *Mlh1* allele harboring promoter methylation in both spleen and jejunum of *Mlh1*^*+/−*^ tissues (**Supplementary fig. 5B**). Unlike *Mlh1*^*+/−*^ mice, none of *Mlh1*^*+/+*^ mice (irrespective of age) (n= 3, 5 and 4 for 1-, 4- and 12-month time point) showed *Mlh1* promoter methylation in jejunum nor in spleen. In contrast, all the *Mlh1*^*−/−*^ mice (n= 3, 3 and 2 for 1-, 4-and 12-month time point) showed *Mlh1* promoter methylation in both the tissues (**Fig. 4B and Supplementary fig. 5A)**.

Further, to investigate if other tissues (except spleen and jejunum) also show *Mlh1* promoter methylation in the *Mlh1*^*+/−*^ mice with *Mlh1* promoter methylation in jejuna, we tested the *Mlh1* promoter methylation status in brain, liver and kidney of these mice (n=3 and 3 for 4- and 12-month time point, respectively). All of these tissues also showed *Mlh1* promoter methylation (**Supplementary fig. 5C)**.

Unlike, jejunum, where *Mlh1* promoter methylation status affected the MLH1 mRNA and protein expression levels, other *Mlh1*^*+/−*^ tissues tested (brain, liver, kidney, spleen), irrelevant of the *Mlh1* promoter methylation status had expected levels of MLH1 expression (i.e. 50% of *Mlh1*^*+/+*^tissue) (**Fig. 3B and Supplementary fig. 5D**).

To investigate the genetic mechanism behind MLH1 depletion we performed LOH assay across the *Mlh1* gene using LOH-PCR and RFLP assay (**Fig. 4A**). LOH was only observed in jejunum of the outlier 4-month old *Mlh1*^*+/−*^ mouse (**Fig. 4C and Fig. 4D**). All other 4- and 12-month old *Mlh1*^*+/−*^ jejuna (n=21) retained their wildtype allele, i.e. no LOH had occurred. In addition, we also tested LOH in spleen of the outlier 4-month old *Mlh1*^*+/−*^ mouse, and found that the wildtype allele was still retained.

### MSI correlates with MLH1 protein level in jejunum of Mlh1^*+/−*^ mice

To explore the tissue- and age-specific relationship between deletions at mononucleotide repeats and MLH1 protein expression, we performed linear regression analysis. In *Mlh1*^*+/−*^ jejuna, deletions were inversely correlated with MLH1 protein expression at both time points (**Fig. 5A**). Further, based on this correlation, at both the time points, *Mlh1*^*+/−*^ mice could be divided into two sub-groups, henceforth termed as sub-grp.1 and sub-grp.2, and defined by comparatively higher MLH1 protein levels and lower deletions at mononucleotide repeats, versus low MLH1 protein levels and higher deletions in jejunum, respectively.

**Fig. 5.**
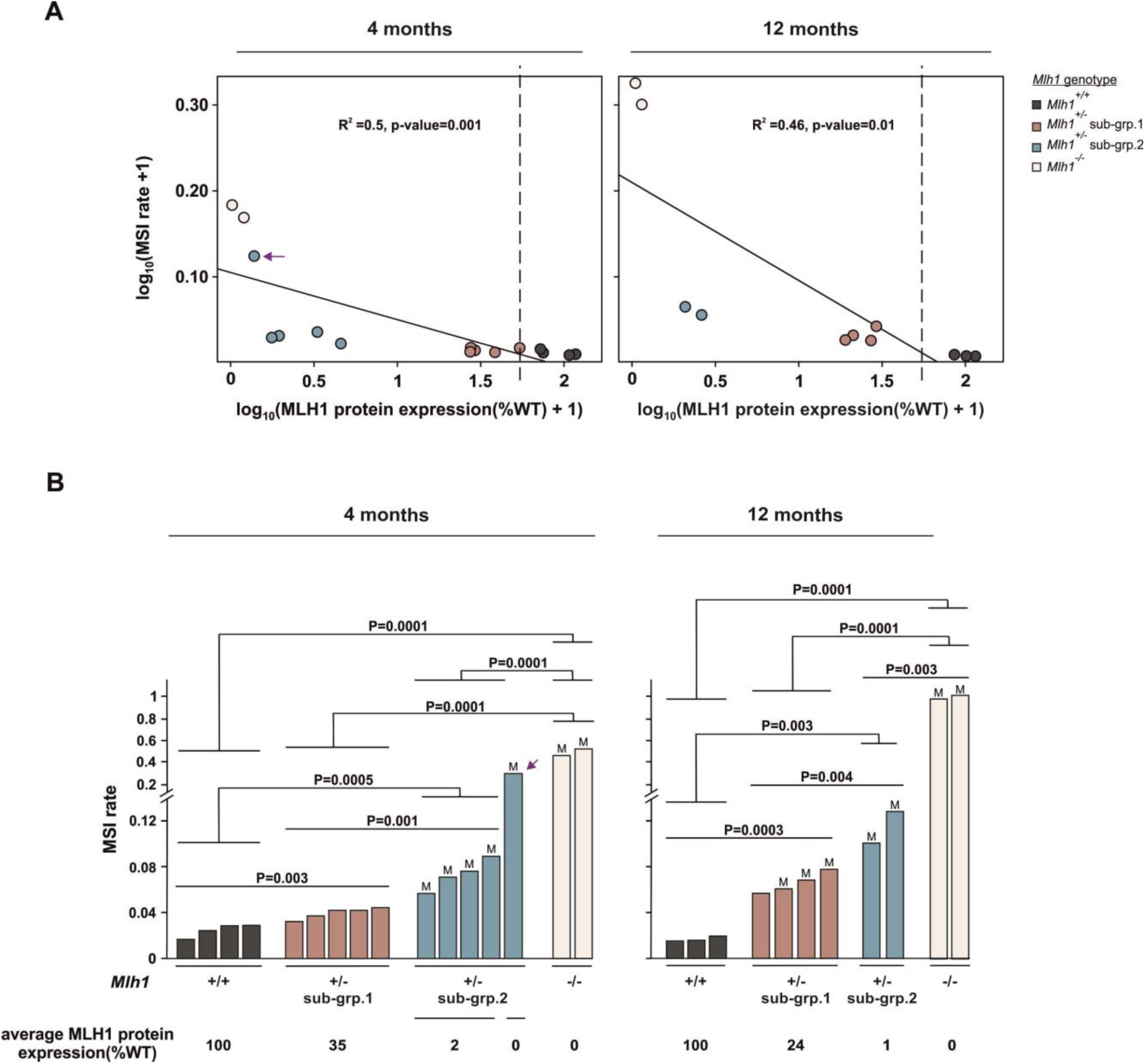
Deletions at mononucleotide repeats and MLH1 protein levels inversely correlate in jejunum of *Mlh1^+/−^* mice. (**A**) Linear regression fitting curve of log_10_(MSI rate+1) ∼ log_10_(MLH1 protein expression(%WT)+1)) at 4- and 12-month time point. Dashed vertical line marks 50% MLH1 expression level. (**B**) Comparison of deletions between the two *Mlh1*^*+/−*^ sub-groups. The arrow indicates the outlier *Mlh1*^*+/−*^ mouse. In **(B)** “M” indicates those *Mlh1*^*+/−*^ jejunum samples where *Mlh1* promoter methylation was detected.

At the 4-month time point, sub-grp.1 showed close-to-expected (approximately 50% of wildtype) average MLH1 protein levels (average 35%, range: 27%-53%) and 1.6-fold higher deletions compared to *Mlh1*^*+/+*^ jejuna. Sub-grp.2 had very low MLH1 protein levels (average: 2%, range: 0%-4%) and displayed 3-fold increase in deletions compared to *Mlh1*^*+/+*^ jejuna (**Fig. 5B**). At the 12-month time point, sub-grp.1 showed lower-than-expected average MLH1 expression (average 24%, range: 18%-29%) and 4-fold increase in deletions compared to *Mlh1*^*+/+*^ jejuna. Sub-grp.2, similarly as observed at 4 months, showed very low MLH1 protein levels (average: 1%, and range: 1%-2%) and showed 7-fold increase in deletions compared to *Mlh1*^*+/+*^ jejuna (**Fig. 5B**). The *Mlh1*^*+/−*^ outlier mouse (indicated by arrow in **Fig. 5A** and **Fig. 5B**) had no detectable MLH1 protein in its jejunum, accompanied by 14-fold increase in deletions compared to age-matched *Mlh1*^*+/+*^ jejuna. Irrespective of age, all *Mlh1*^*+/−*^ spleens showed the expected approximately 50% MLH1 protein level, and no increase in deletions compared to *Mlh1*^*+/+*^ spleens except spleen of outlier *Mlh1*^*+/−*^mouse which had double the deletions compared to age-matched *Mlh1*^*+/−*^ spleens despite having normal protein expression level (**Supplementary fig. 9**).

## Discussion

*Mlh1* mice [7, 22] are powerful model to comprehensively study MSI, and to dissect molecular mechanisms contributing to the MSI phenotype. In line with previous studies [29, 39, 40], we observed mononucleotide repeats to be more unstable than dinucleotide repeats. The key to detecting subtle cellular phenotypes with pre-malignant potential, such as low-level MSI, is use of sufficiently sensitive molecular read-outs. The SM-PCR assay, when applied to unstable microsatellite loci, can detect MSI levels as low as 1%, whereas standard PCR has a detection limit of 20-25% MSI [41]. Analyzing less unstable microsatellite markers by conventional PCR may result even in failure to detect MSI in *Mlh1*^*+/−*^ tumors [36].

We observed increase in MSI with age in *Mlh1*^*+/−*^ mice, however, 42% of young (4-months) *Mlh1*^*+/−*^ mice showed similar or even more (case of outlier 4-month *Mlh1*^*+/−*^ mice) MSI than the old (12-months) mice. Further, we observed variation in jejuna-specific MSI in *Mlh1*^*+/−*^ mice of same age group. These observation i.e. individuals showing elevated MSI at young age, and inter-individual variation in MSI is similar to that reported in peripheral blood leukocyte DNA of LS patients [15]. In addition, in our study, 1-month-old *Mlh1*^*+/−*^ mice also showed jejunum-specific inter-individual variation in Mlh1 mRNA levels, and lower than expected Mlh1 mRNA levels, suggesting that the cellular changes (including MSI) may occur even earlier.

LS associated colorectal cancers are MMR-deficient and show high-level MSI, which led to the notion that two hits for MMR genes are required to instigate the MSI phenotype [9, 11]. However, recent evidence that LS-associated adenomas retain (some) MMR function suggests that the second hit may arise later in the multi-step tumorigenesis than previously appreciated [42], making the idea of pre-tumorigenic MMR haploinsufficiency more plausible. We demonstrate here that MSI is present in normal jejunum of *Mlh1*^*+/−*^ mice with expected (i.e. 50% of wildtype) MLH1 levels, providing evidence for MMR haploinsufficiency.

Surprisingly, all *Mlh1*^*+/−*^mice harboring *Mlh1* promoter methylation in jejunum also showed *Mlh1* promoter methylation in other tissues analyzed. We observed *Mlh1* promoter methylation in tissues originating from all three germ layers: ectoderm-brain, mesoderm-kidney and spleen, endoderm-jejunum and liver, indicating that the epigenetic modification occurred during (or even earlier) than early-embryonic development in these mice. Our study reports this observation for the first time in MMR mouse model, which very well mimics the observation (soma-wide *Mlh1* promoter methylation) in LS (suspected) patients (individuals with similar clinicpathological phenotype as LS, but no germline MMR mutations) [43-49]. Upon investigating the allele specificity of the *Mlh1* promoter methylation, we found out that the *Mlh1* promoter methylation occurred in *Mlh1*^*−/−*^allele of *Mlh1*^*+/−*^ DNA. Promoter methylation of the *Mlh1*^*−/−*^allele tends to influence the MLH1 protein expression in a tissue-specific manner (specifically in intestine), and was observed only in fraction of *Mlh1*^*+/−*^mice with the soma-wide *Mlh1* promoter methylation. As we used targeted approach to test the promoter methylation, we cannot negate the possibility of methylation of other CpG sites of the *Mlh1* promoter region of *Mlh1* ^*+/+*^allele leading to tissue-specific MLH1 depletion. Additionally, heterozygosity of *Mlh1* is known to provoke promoter methylation and as consequence lower expression of multiple tumor suppressor genes in normal GI tract affecting crucial signaling pathways [25]. It is likely, that the lower tissue-specific *Mlh1* expression we observed is consequence of altered signaling pathways in *Mlh1*^*+/−*^ intestine. However, given that we observed a strong correlation of site-specific promoter methylation in *Mlh1*^*−/−*^ allele and elevated MSI, we propose site-specific promoter methylation assay around transcription start site of *Mlh1* as a good reporter of tissue-specific per-neoplastic MSI.

In human, LS individuals are most susceptible to colorectal and endometrium cancer [43-48]. In *Mlh1*^*+/−*^ mice, though tumor incidence is very low, GI tract tumors, and non-GI tract tumors namely lymphoma, cervical squamous cell carcinoma and lung bronchio-alveolar carcinoma has been reported [20]. In both cases (human and mice), the tumor spectra due to defective MMR comprises of proliferative-tissues or -cell types (hematopoietic system). In addition, with our observation of tissue specific pre-tumorigenic cellular events (including MSI) in intestine (a tissue with high proliferation rate) of *Mlh1^+/−^* mice, a plausible assumption is that these tissues (with high proliferation rate) are more vulnerable to pre-tumorigenic MMR associated abnormalities (making them susceptible to tumorigenesis) due to higher likelihood of accumulating replication errors in every cell division. Additionally, MMR also plays role in cell cycle arrest and apoptosis [50], in case of faulty MMR these functions are inhibited and replication errors accumulates in every subsequent cell divisions. Our observation of pre-tumorigenic cellular events in jejunum may also prevail in other tissues/cell types with high turnover rate (such as endometrium) making them venerable to MMR-associated tumorigenesis in LS condition, which we propose as a future line of investigation.

Interestingly, we also observed inter-individual variation in MSH2 protein levels in *Mlh1*^*+/−*^ jejuna, with significant fraction of *Mlh1*^*+/−*^ jejuna expressing lower MSH2 than the *Mlh1*^*−/−*^ jejuna. Further, we saw a significant decrease in MSH2 protein levels with *Mlh1* gene dosage in both jejunum and spleen in our western blot assay, while Msh2 mRNA levels were unaltered. These observations suggest a protein-level dependency in stability of the two heteroduplex MutS homolog (MSH2-MSH6) and MutL homolog (MLH1-PMS2), and that *Mlh1*^*+/−*^ jejuna are more liable for MMR protein destabilization than *Mlh1*^*−/−*^ jejuna. However, with IHC analysis, we saw no substantial difference in MSH2 levels in *Mlh1*^*+/+*^, *Mlh1*^*+/−*^ and *Mlh1*^*−/−*^ jejuna (except one *Mlh1*^*+/−*^ jejuna). Similar histopathological observation (i.e. MSH2 not affected by MLH1 levels) is typical in *Mlh1*-deficient MSI-positive CRC tumors in human [51]. These observations indicate western blot assays are more sensitive in detecting wide range of protein expression variation than IHC.

MSI reports on global MMR-dependent genome instability, but MSI *per se* does not necessarily lead to tumorigenesis [20]. Tumorigenesis initiates when MSI occurs in microsatellite-containing genes, giving the cells selective advantage and making them malignant. These genes include tumor suppressors and oncogenes involved in different regulatory pathways, including cell proliferation, cell cycle, apoptosis, and DNA repair [52, 53]. As mismatches within genes (particularly exons) enjoy more efficient MMR compared to intergenic regions [54], it is unlikely that MSI-target genes would have early mutation accumulation, which could alter gene function. We did not observe any tumors in *Mlh1*^*+/−*^ jejunum, although this observation was limited to mice that were sacrificed to harvest tissues (We did not examine post-mortem the 30% *Mlh1*^*+/−*^ mice which died in our care; **Supplementary fig. 10**). The MSI detected in intergenic/intronic microsatellites in normal *Mlh1*^*+/−*^ jejuna reports on early-stage genomic instability prior to tumorigenesis. The 4-month-old outlier *Mlh1*^*+/−*^ mouse which had comparable deletions in its jejunum as age-matched *Mlh1*^*−/−*^ mice would presumably have had a higher chance of accumulating mutations in MSI-target genes that, over time, may have triggered tumorigenesis in the GI tract of this animal. Also, it seems likely that many, if not all, of those *Mlh1*^*+/−*^ mice for which the cause of death is unknown (**Supplementary fig. 10**), succumbed to MMR-associated tumors.

Ours is the first systematic study that establishes the relationship between MLH1 levels and MSI in a variety of normal tissues. MSI in non-neoplastic mucosa [16] and crypt cells [55] has been reported in LS patient samples that likely were MMR-deficient. We demonstrate here that MSI is detectable – albeit at a relatively low level – in *Mlh1*^*+/−*^ jejunum that still retains the expected 50% MLH1 protein level, implicating MMR haploinsufficiency in normal, tumor-free *Mlh1*^*+/−*^ intestine. High MSI only ensued upon substantial reduction of MMR protein levels. Thus, MMR function appears to rely on a critical threshold level of MMR proteins, and is independent of age. Importantly, MLH1 expression aberration was tissue-specific, which was observed only in jejunum and not in other tissues assayed.

Given that maintenance of genome stability is essential for all cells, it has been puzzling why germline mutations of key DNA repair pathways (mismatch repair, homologous recombination) give rise to cancer only in certain tissues. The tissue-specific haploinsufficiency of *Mlh1*, induced by promoter methylation, now provides an explanation for vulnerability of the GI tract to MMR-associated cancer. Heterozygosity of homologous recombination genes, namely *BRCA1* and *PALB2*, also confers subtle genome instability phenotypes that are detectable in pre-malignant cells of mutation carriers, provided that sufficiently sensitive assays are used [56, 57]. We propose that tissue-specific decrease in protein levels is an important factor in determining which organs are cancer-prone in heterozygous DNA repair mutation carriers.

## Author contributions

K.S.S. designed and performed the experiments, analyzed the data, interpreted the results and wrote the manuscript. L.K. conceived and designed the study, interpreted the data, reviewed and edited the manuscript, acquired funding and supervised K.S.S. M.M.T. assisted in tissue collection, MSI assay, qPCR experiments, and genotyping. E.A. gave advice on calculation of MSI rate and edited the manuscript.

## Acknowledgments

We are grateful to Erika Gucciardo, Manuela Tumiati, Päivi Peltomäki, Marjaana Pussila and Saara Ollila for critical reading of the manuscript, to Elina A Pietilä and Joonas Jukonen for their advice for western-blotting experiments, and to members of the Kauppi lab for constructive comments on the figures. We extend our sincere gratitude to the staff of the following core facilities at University of Helsinki: Laboratory Animal Center, Tissue preparation and histochemistry unit, Genome Biology Unit, and FIMM sequencing unit.

## Supplementary figures

**Supplementary fig. 1.**
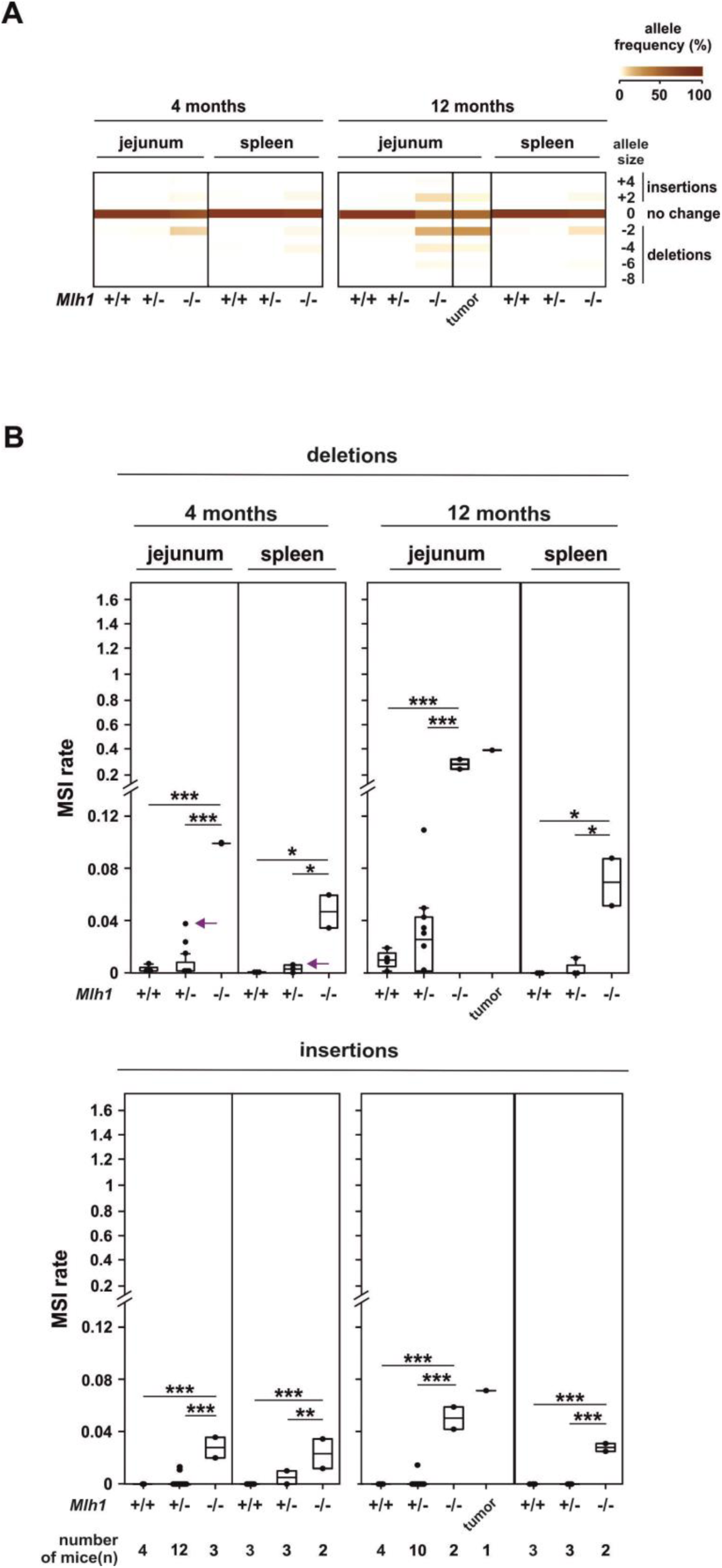
Single-molecule MSI analysis at dinucleotide D14Mit15. (**A**) Heat map of the % of different-sized alleles at microsatellite D14Mit15. (**B**) MSI in normal jejunum and spleen. Indicated are significant p-values for the deletions (top panel) and insertions (bottom panel) comparisons. The arrow indicates the outlier *Mlh1*^*+/−*^ mouse.

**Supplementary fig. 2.**
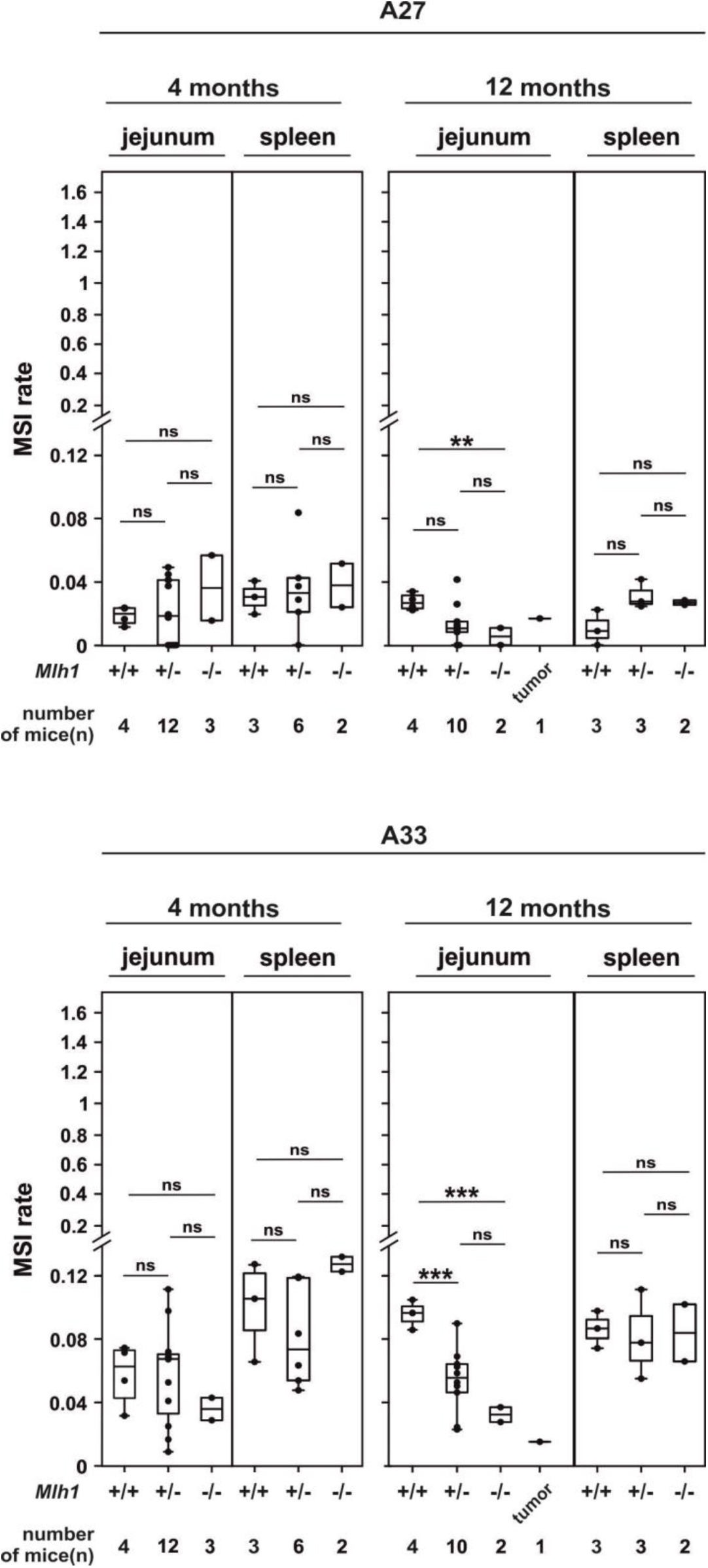
Insertions at mononucleotide repeats in jejunum and spleen. Top and bottom panel shows insertions at A27 and A33, respectively.

**Supplementary fig. 3.**
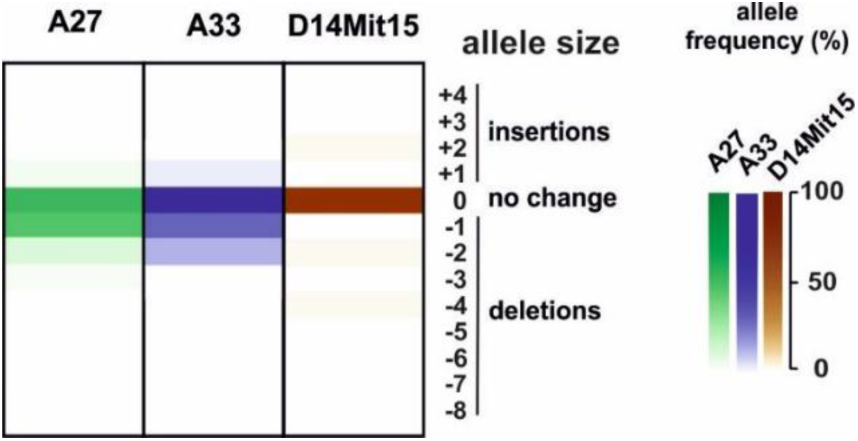
MSI in the outlier *Mlh1*^*+/−*^ mouse. Heat map of % of allele sizes for microsatellites A27, A33 and D14Mit15 observed in jejunum of outlier *Mlh1*^*+/−*^ mouse with elevated deletions at mononucleotide repeats.

**Supplementary fig. 4.**
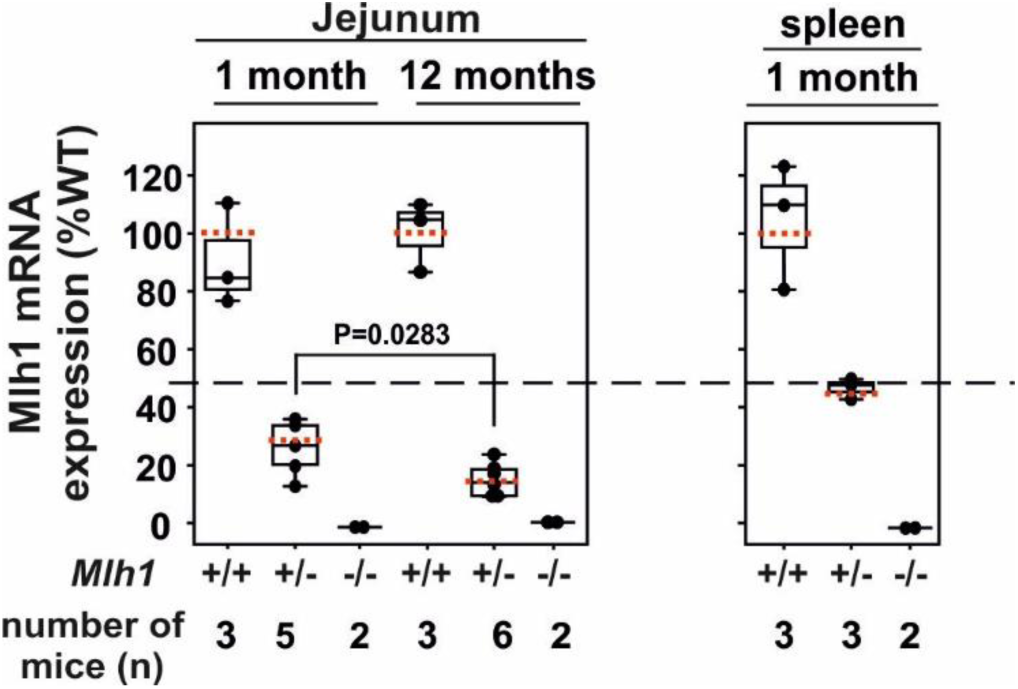
Comparison of Mlh1 mRNA expression in 1- and 12-month old mice. The dashed horizontal line indicates 50% Mlh1 mRNA expression. The red dotted line across each box-plot shows the average value for each genotype group. Indicated is the only statistically significant p-value for within-genotype comparisons of Mlh1 mRNA levels in the jejunum in 1-vs. 12-month old mice.

**Supplementary fig. 5.**
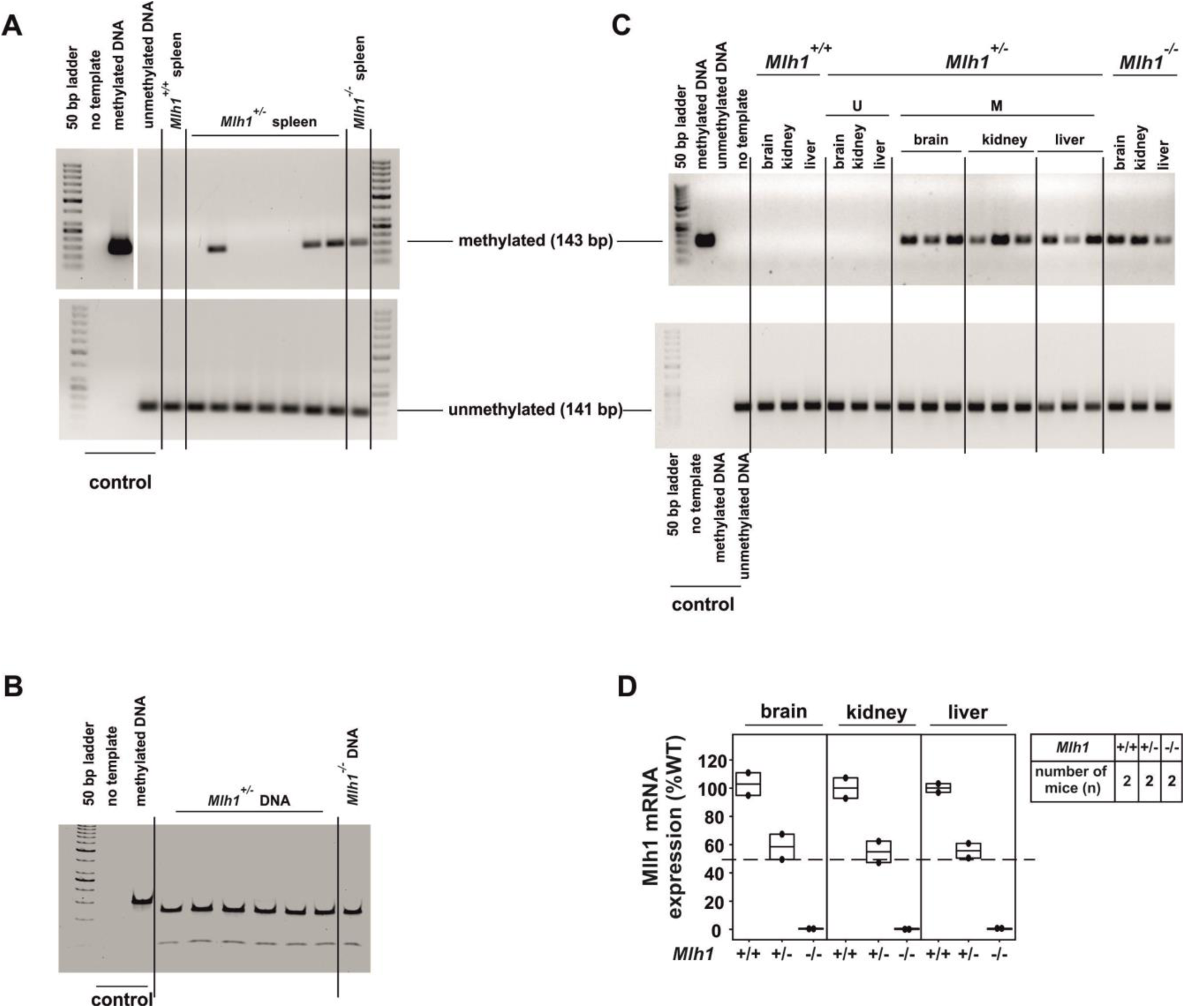
*Mlh1* promoter methylation and *Mlh1* gene expression analysis in various tissues of *Mlh1+/−* mice. Representative gel images of (A) MSP assay for spleen, (B) RFLP assay on methylated-PCR-product fraction of the MSP assay, (C) MSP assay for brain, kidney and liver, respectively (“U” and “M” indicates *Mlh1*^*+/−*^ mice with and without *Mlh1* promoter methylation, respectively. (D) Mlh1 mRNA expression in brain, kidney and liver of *Mlh1*^*+/−*^ mice with *Mlh1* promoter methylation show approximately 50% expression level compared to respective wildtype tissue. The dashed horizontal line indicates 50% of wild-type Mlh1 mRNA expression. MLH1 expression data for spleen is shown in **Fig. 3A** and **Fig. 3B**.

**Supplementary fig. 6.**
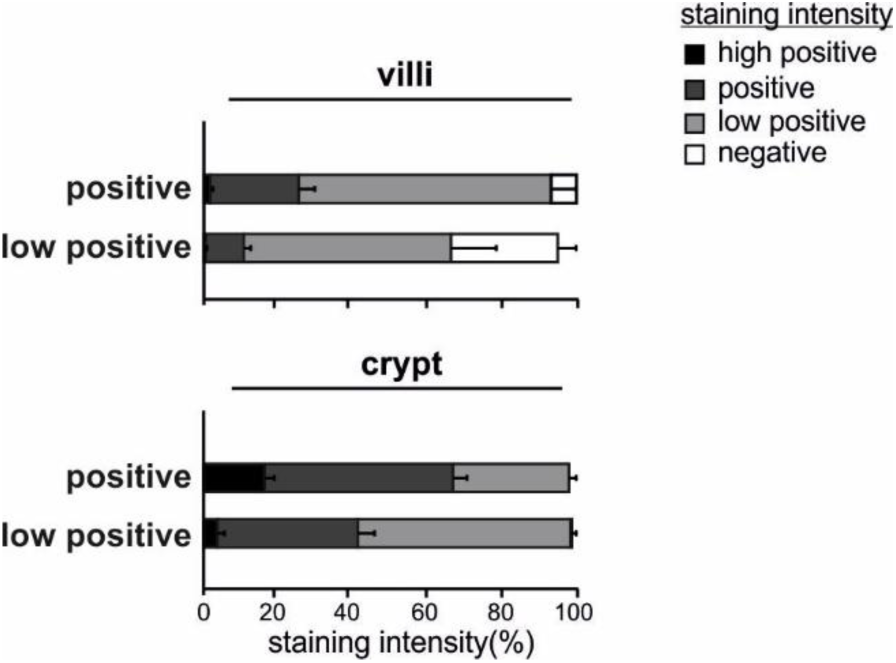
Comparison of MLH1 staining intensity in crypts and villi of MLH1-positive and MLH1-low positive *Mlh1*^*+/−*^ jejunum samples. The stackbar shows mean + SD.

**Supplementary fig. 7.**
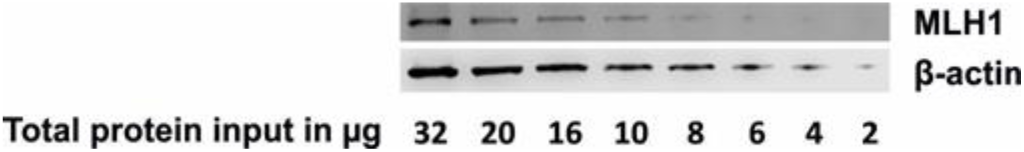
Sensitivity of western blot assay. A dilution series of protein was run to determine sensitivity of the western blot assay. Infra-red signal was not detectable below 4 µg of MLH1 protein.

**Supplementary fig. 8.**
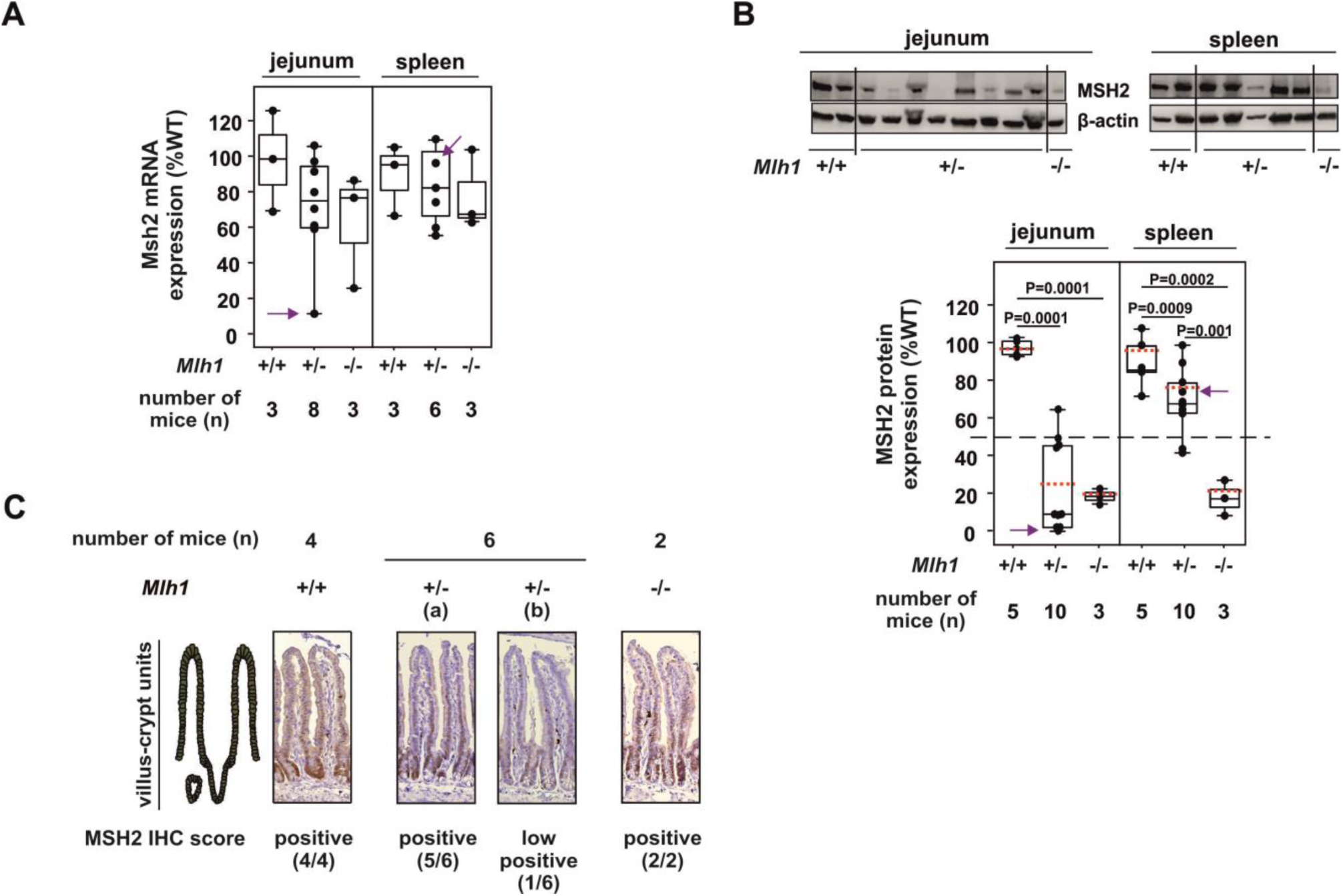
*Msh2* gene expression analysis in jejunum and spleen of *Mlh1+/−* mice at 4-month time point. (**A**) Msh2 mRNA expression analysis. Arrow indicates data from the outlier *Mlh1*^*+/−*^ mouse with high deletions at mononucleotide repeats. (**B**) and (**C**) MSH2 protein levels analysis. (**B**) Representative image of western blot. Boxplot shows the MSH2 protein expression analysis for western blot. The red-dotted line across each box-plot represents the average value. The dashed-horizontal line across the chart represents the 50% MSH2 protein expression. (**C**) Representative IHC images of jejunum. Middle two IHC images shows a side-by-side comparison of villus-crypt units scored as (a) positive and (b) low positive by IHC profiler. Only significant p-values are showed in **Supplementary fig. 8A** and **Supplementary fig. 8B**.

**Supplementary fig. 9.**
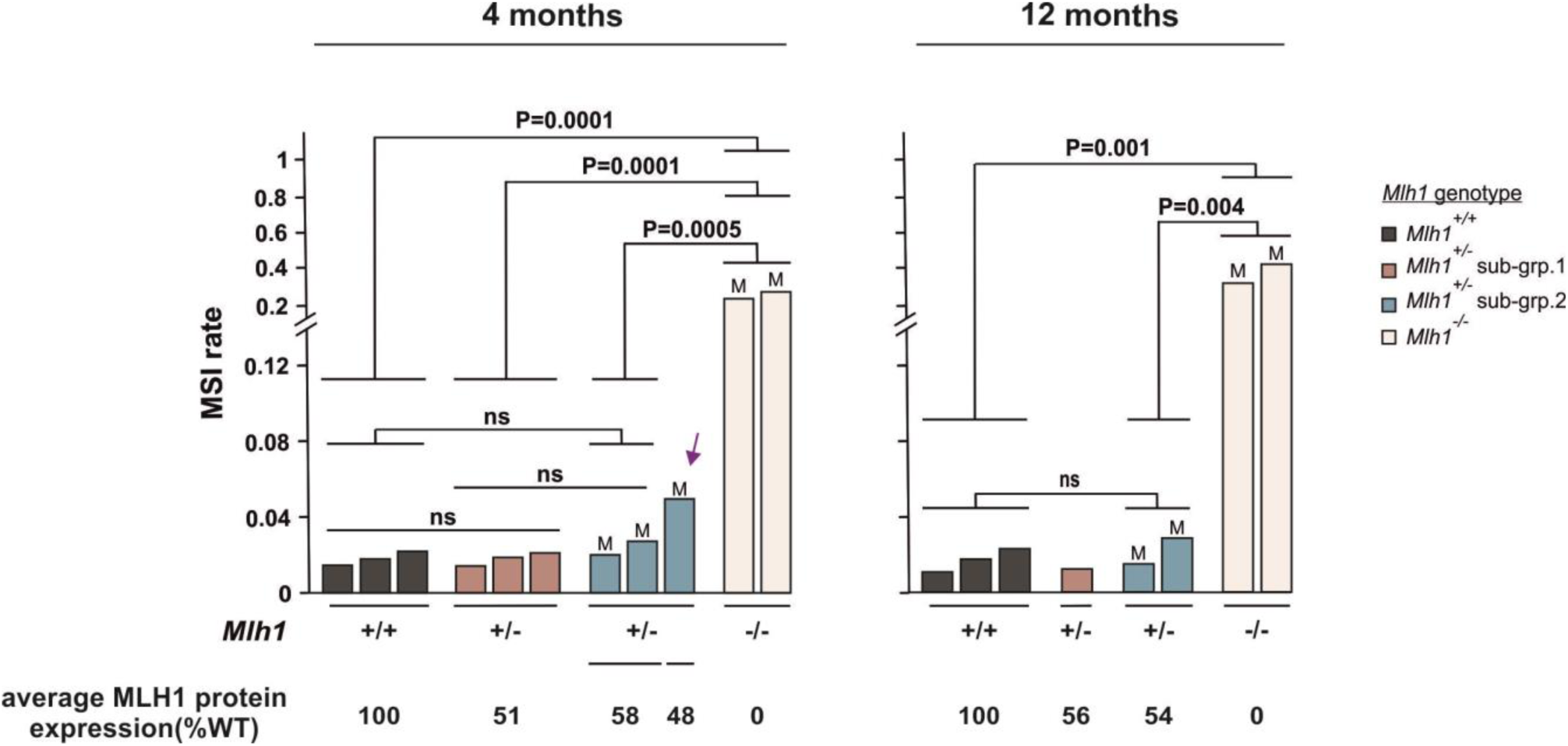
Relationship between deletions and MLH1 protein levels in spleen of *Mlh1+/−* mice. The arrow indicates the outlier *Mlh1*^*+/−*^ mouse. “M” indicates those *Mlh1*^*+/−*^ spleen samples where *Mlh1* promoter methylation was detected.

**Supplementary fig. 10.**
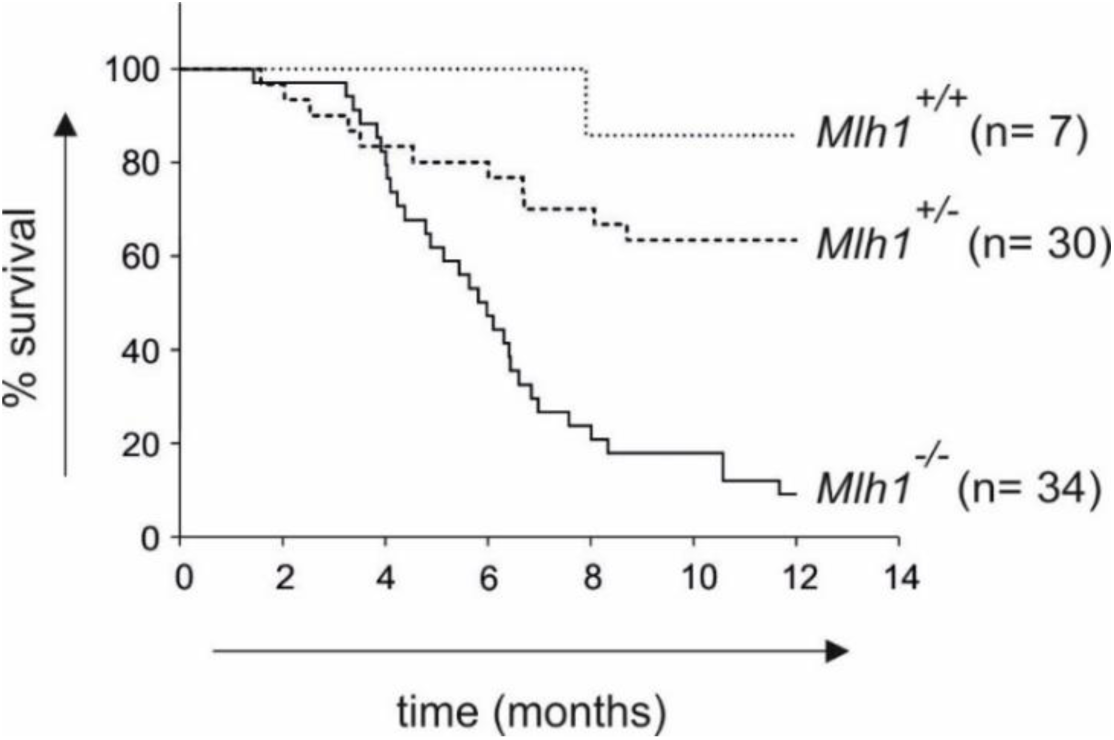
Kaplan-Meier survival curves for *Mlh1* mutant mice.

**Supplementary table 1.**
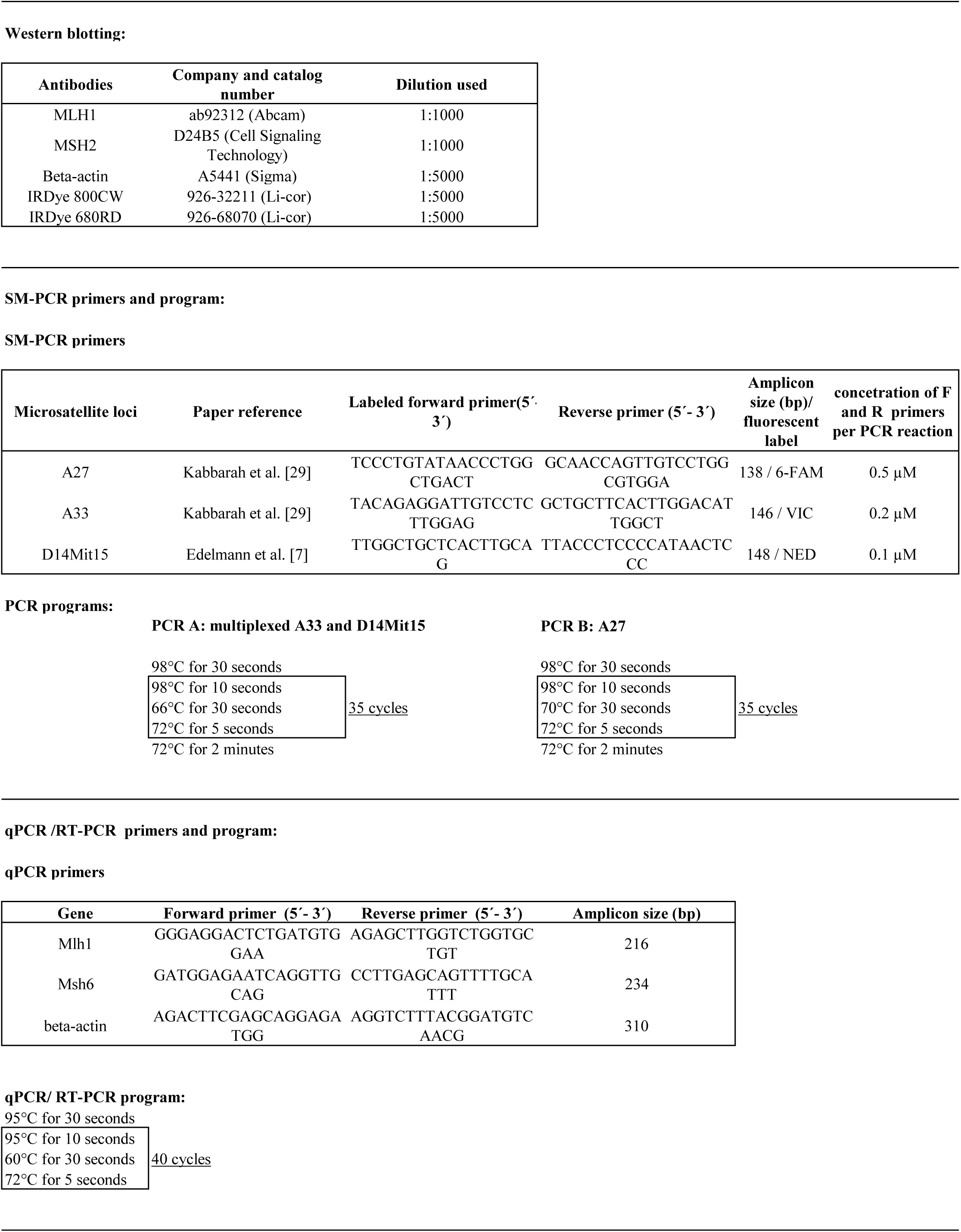
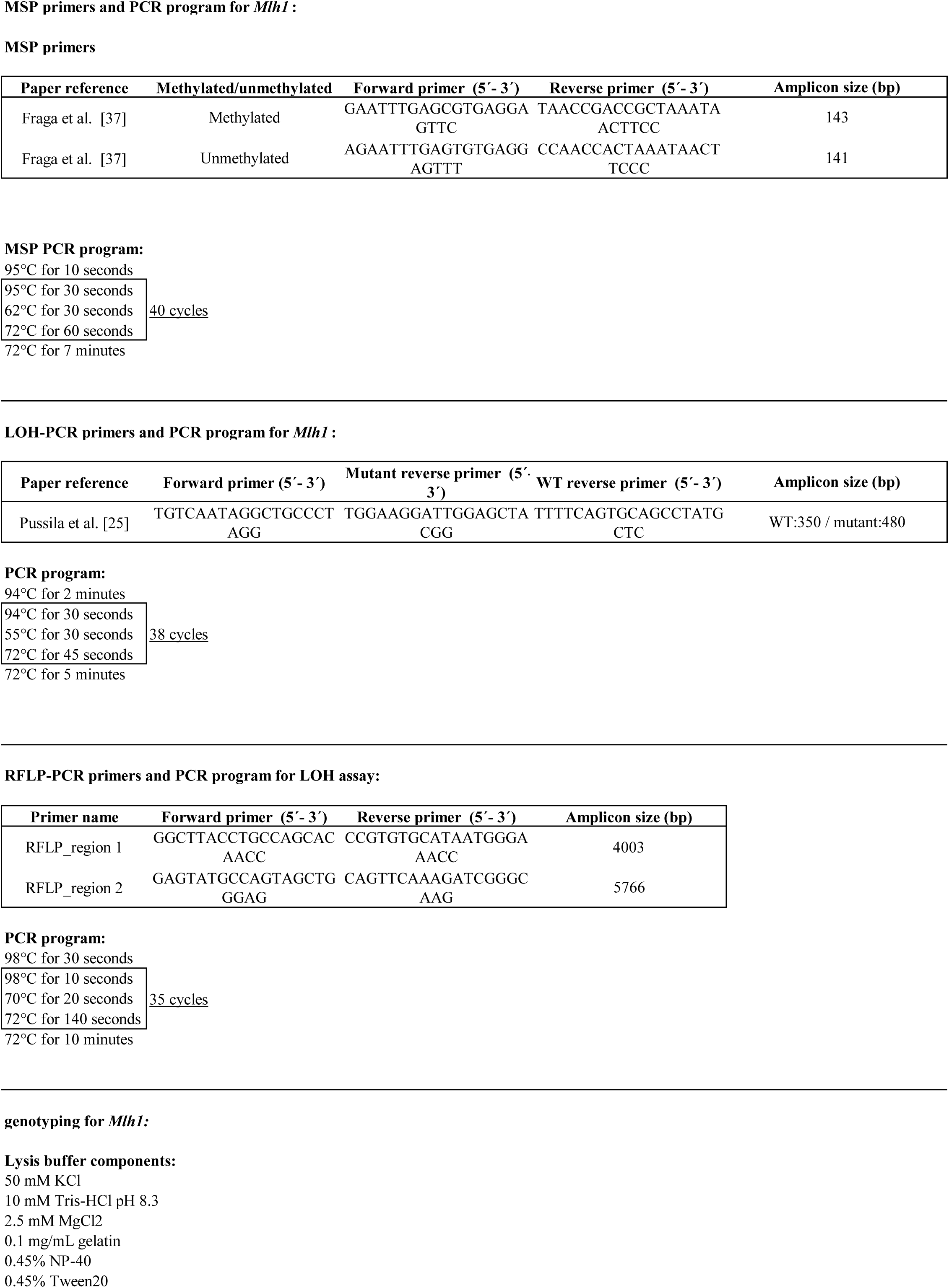
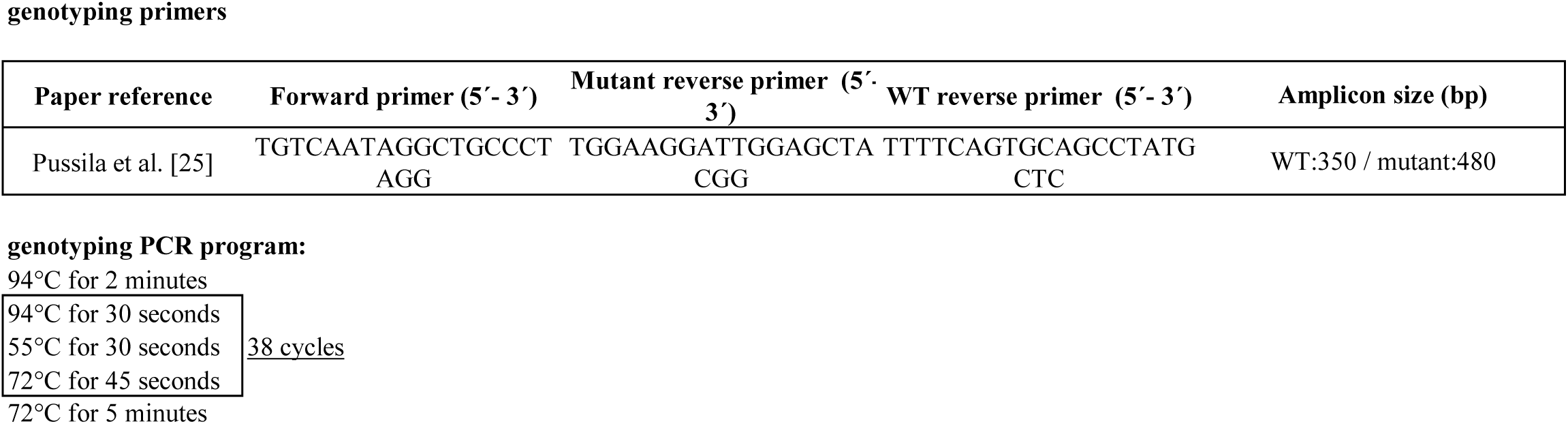

